# FICD acts bi-functionally to AMPylate and de-AMPylate the endoplasmic reticulum chaperone BiP

**DOI:** 10.1101/071332

**Authors:** Steffen Preissler, Claudia Rato, Luke Perera, Vladimir Saudek, David Ron

**Author notes:** Equal contribution (Steffen Preissler, Claudia Rato). Corresponding authors: David Ron and Steffen Preissler, Cambridge Institute for Medical Research, University of Cambridge, Cambridge Biomedical Campus, Wellcome Trust/MRC Building, Hills Road, Cambridge CB2 0XY, United Kingdom.

## Abstract

**Significance statement:** Some 25 years ago it was discovered that the activity of the ER chaperone BiP is regulated by covalent modification, the nature of which, AMPylation (not ADPribosylation, as had long been thought) and the enzyme responsible, FICD, have only recently been identified. Genetic inactivation of FICD and *in vitro* studies of the purified enzyme and substrate have done much to clarify the biochemical consequences of the modification and its underlying logic: As ER stress wanes, FICD uses ATP to AMPylate Thr518 of BiP locking BiP in a relatively inactive conformation. As ER stress levels re-mount the cells draw on this pool of inactive chaperone, which is de-AMPylated and restored to its fully active state.

Here we report on **the identity of the de-AMPylating enzyme** - and with it on the surprising finding that both AMPylation and de-AMPylation of BiP are carried out by the same polypeptide (FICD) using the same active site, both in vivo and in vitro. Analysis of the reaction products reveals that de-AMPylation does not involve trivial concentration-dependent micro-reversibility of an enzymatic reaction, but rather a switch in the active site of FICD that facilitates two antagonistic thermodynamically favored reactions.

Surprisingly **BiP de-AMPylation** (not AMPylation) is the **default activity of FICD**. The side-chain of **a single regulatory residue, E234, toggles the enzyme** between de-AMPylation and AMPylation in vitro. Our studies thereby uncover **an active mechanism** that must exist in the ER for coupling waning levels of unfolded protein stress to the conversion of FICD from its default de-AMPylation mode to BiP AMPylation. Whilst the details of this active switch remain to be discovered, we are able to suggest a plausible mechanism by which it may come about.

Identification of the enzyme that de-modifies BiP to reactivate it will be of interest to cell biologists, whereas the novel features of FICD as a dual-functioning enzyme with a single bi-functional active site will be of broad interest to enzymologists and molecular biologists.

**Abstract:** Protein folding homeostasis in the endoplasmic reticulum (ER) is defended by an unfolded protein response (UPR) that matches ER chaperone capacity to the burden of unfolded proteins. As levels of unfolded proteins decline, a metazoan-specific FIC-domain containing ER-localized enzyme, FICD/HYPE, rapidly inactivates the major ER chaperone BiP by AMPylating T518. Here it is shown that the single catalytic domain of FICD can also release the attached AMP, restoring functionality to BiP. Consistent with a role for endogenous FICD in de-AMPylating BiP, *FICD*^*−/−*^ cells are hypersensitive to introduction of a constitutively AMPylating, de-AMPylation defective mutant FICD. These opposing activities hinge on a regulatory residue, E234, whose default state renders FICD a constitutive de-AMPylase in vitro. The location of E234 on a conserved regulatory helix and the mutually antagonistic activities of FICD in vivo, suggest a mechanism whereby fluctuating unfolded protein load actively switches FICD from a de-AMPylase to an AMPylase.

## Introduction

The balance of chaperones and unfolded proteins in the endoplasmic reticulum (ER) is important to the functionality and health of all secretory cells and affects the outcome of diverse protein misfolding and aging-related diseases (Balch et al., 2008; Wang and Kaufman, 2016). Conserved transcriptional and translational mechanisms, operative on a timescale of hours, match ER folding capacity to client protein abundance in all eukaryotes (Walter and Ron, 2011). In metazoans this unfolded protein response (UPR) is complemented by processes that rapidly inactivate and reactivate the ER-localized Hsp70 chaperone BiP to match fluctuating levels of unfolded proteins during the inherent delay of the UPR.

Two processes are known to contribute to this short-term post-transcriptional buffering. Firstly, client protein binding is in competition with rapid self-binding of BiP to form oligomers that serve as a pool of recruitable inactive chaperone (Chevalier et al., 1998; Freiden et al., 1992; Preissler et al., 2015a). Secondly, an enzymatically-mediated inactivating covalent modification of BiP, which is conspicuous when unfolded proteins are scarce (Chambers et al., 2012; Laitusis et al., 1999) functions alongside mass-action mediated oligomerization to match BiP activity to client protein load. Long believed to be ADP-ribosylation (Carlsson and Lazarides, 1983; Freiden et al., 1992), this modification is now known to be AMPylation (Ham et al., 2014; Preissler et al., 2015b; Sanyal et al., 2015); the covalent attachment of adenosine mono phosphate, via a phosphodiester bond to the hydroxyl side chain of a residue in the target protein (also known as adenylylation).

The ER-localized FIC- (<u>f</u>ilamentation <u>i</u>nduced by <u>c</u>yclic AMP) domain containing protein, FICD/HYPE (Worby et al., 2009), uses ATP to AMPylate BiP both in vitro and in vivo (Broncel et al., 2015; Ham et al., 2014; Preissler et al., 2015b; Sanyal et al., 2015). In cultured mammalian cells, deletion of the *FICD* gene abolishes all evidence for BiP modification, which is otherwise observed at high stoichiometry on threonine 518. AMPylated BiP (BiP^T518-AMP^) is only weakly stimulated by Jdomain proteins and the modified chaperone is locked in a relatively inert state (Preissler et al., 2015b). Enforced expression of a constitutive AMPylating FICD mutant results in high levels of ER stress – most likely a consequence of BiP inactivation. These genetic and biochemical findings point to FICD as being both necessary and sufficient for BiP AMPylation observed when the burden of unfolded ER proteins is low.

As unfolded proteins accumulate, pre-existing AMPylated BiP is rapidly converted to the active de-AMPylated state (Chambers et al., 2012; Ham et al., 2014; Laitusis et al., 1999; Preissler et al., 2015a; Preissler et al., 2015b), indicating that AMPylation is a reversible modification that contributes to the balance between clients and chaperone in the ER. This paper addresses the hitherto mysterious process by which the phosphodiester bond between AMP and the hydroxyl side chain of BiP's T518 is broken and full BiP chaperone activity restored.

## Results

AMPylated BiP, detected by its characteristic mobility on native-PAGE or in isoelectric focusing gels, is readily observed upon inhibition of protein synthesis in wildtype but not *FICD*^−/−^ cells (Figure 1). Overexpression of FICD fails to restore AMPylation to *FICD*^−/−^ cells, suggesting that elaborating a pool of AMPylated BiP is sensitive to FICD dosage. By contrast, the hyperactive allele FICD^E234G^ readily restored AMPylated BiP in *FICD*^−/−^cells [Figure 1, lanes 4 & 5 and (Preissler et al., 2015b, figure 2D therein)]. This finding suggested the possibility that levels of AMPylated BiP are biphasically-related to the concentration of wildtype (but not mutant FICD^E234G^) and that in addition to AMPylating BiP, wildtype FICD also has a role in undoing the modification; a role that dominates in overexpressionprone *trans*-rescue experiments.

**Figure 1.**
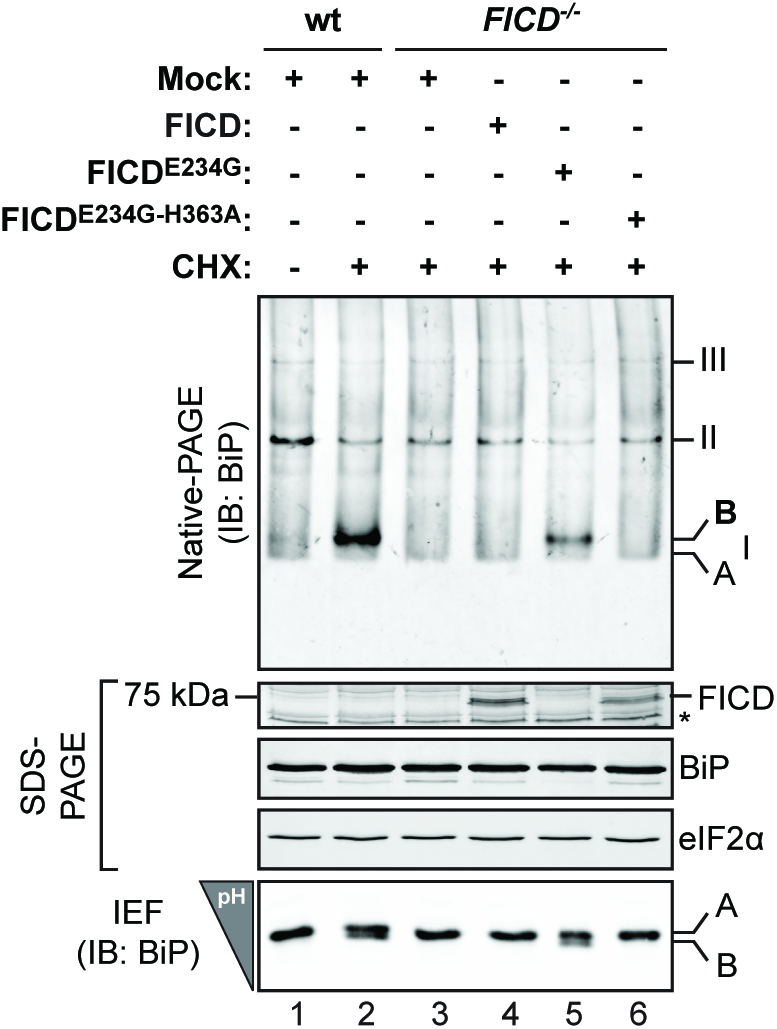
Introduction of wild type FICD into FICD deficient cells fails to restore BiP AMPylation. Immunoblot of endogenous BiP resolved by native gel electrophoresis. Wildtype (wt) and FICD deficient (-/-) CHO-K1 cells were transfected with plasmids encoding the indicated FICD derivatives and exposed to cycloheximide (CHX; 3 hours; 100 µg/mL) to promote AMPylated BiP. The major species visible on the native gel are numbered by order of descending mobility (I-III). Species I represents monomeric BiP whereas species II and III represent BiP oligomers. The monomeric AMPylated ‘B’ form induced by CHX treatment and the ‘A’ form detectable in untreated cells are marked. Immunoblots of the same samples resolved by SDS-PAGE report on FICD, total BiP and total eIF2a (which also serves as a loading control) and the acidic AMPylated ‘B’ form of BiP on the isoelectric focusing gel (IEF). The asterisk indicates a band of unknown identity. Data representative of four independent experiments are shown (*n* = 4). Note the lack of AMPylated BiP in the lysate of *FICD*^−/−^ cells expressing high levels of wildtype FICD.

**Figure 2.**
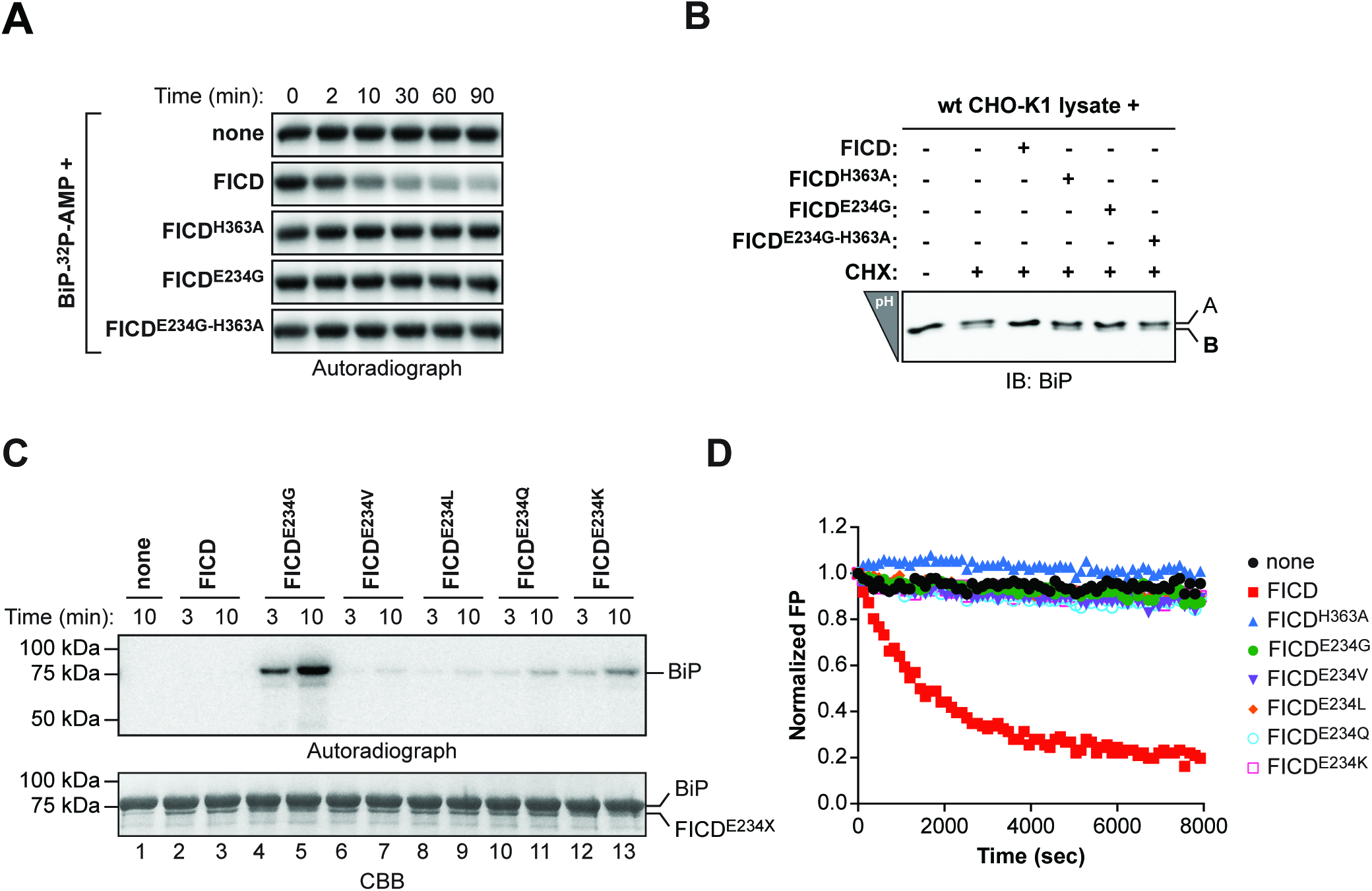
FICD de-AMPylates BiP in vitro. (**A**) Autoradiograph of AMPylated BiP (BiP-^32^P-AMP) separated on an SDS-PAGE gel. Radiolabeled BiP (free of contaminating AMPylating enzyme) was exposed for the indicated time to pure recombinant wildtype or mutant FICD proteins. (**B**) IEF immunoblots of endogenous BiP from lysates of untreated wildtype CHOK1 cells or cells treated with cycloheximide (CHX; 3 hours; 100 µg/mL). Where indicated, purified wildtype or mutant FICD was added to the lysates and incubated for 15 minutes at 30°C before denaturation in IEF lysis buffer followed by IEF. (**C**) Autoradiograph and Coomassie stain (CBB) of an SDS-PAGE gel of BiP after exposure to wildtype or mutant versions of FICD in presence of α-^32^P-ATP. (**D**) Time-dependent plot of fluorescence polarization (FP) of BiP AMPylated with FAM-labeled AMP (BiP^T518-AMP-FAM^) following exposure to the indicated FICD proteins. The decrease in the FP signal reflects release of the fluorophore from BiP.

Speculation that failure to rescue AMPylation reflects narrow tolerance for FICD dosage effects consequent to opposing actions of a single FICD enzyme is further supported by observations that AMPylation and de-AMPylation of bacterial glutamine synthetase is carried out by two structurally-related nucleotidylyl transferase-like domains on the same polypeptide (Xu et al., 2010). Whilst the active sites of the aforementioned Glutamine Synthetase Adenylyl Transferase (GS-ATase) are not obviously related to FIC, the FIC fold also flexibly catalyzes diverse phospho-transfer reactions (Garcia-Pino et al., 2014; Harms et al., 2016; Roy and Cherfils, 2015). Therefore, it seemed reasonable to examine the ability of FICD to remove the AMP moiety from BiP^T518-AMP^.

The signal arising from the 32P-labeled AMP of purified BiP^T518-AMP^ is very stable, with negligible rates of spontaneous hydrolysis. However, addition of wildtype FICD led to a time-dependent decrease in radiolabel; a feature shared neither by the AMPylation-competent, hyperactive FICD^E234G^ nor by the AMPylation-defective FICD^H363A^ mutant enzyme (Figure 2A). These de-AMPylation reactions were performed in the absence of ATP, precluding re-AMPylation by any residual hyperactive AMPylating FICD^E234G^ (carried over from the preceding AMPylation reaction). The de-AMPylating activity of wildtype FICD was not restricted to recombinant BiP AMPylated *in vitro*, as incubating lysates from cycloheximide-treated cells with pure wildtype FICD also led to disappearance of the acidic AMPylated form of endogenous BiP, as revealed by isoelectric focusing (Figure 2B).

Wildtype FICD's ability to catalyze the removal of AMP from BiP^T518-AMP^ was confirmed kinetically by following the decline in the fluorescence polarization signal arising from BiP AMPylated in vitro with fluorescent ATP-FAM as a substrate (BiP^T518-AMP-FAM^; Figure 2 supplement 1). Importantly, different substitutions of FICD^E234^ that differ widely in their AMPylating activity (Figure 2C) all lacked detectable de-AMPylating activity (Figure 2D), attesting to the dual role of this residue in regulating AMPylation (Bunney et al., 2014; Engel et al., 2012) and in affecting de-AMPylation.

To further characterize the FICD-mediated de-AMPylation reaction its products were analyzed. Comparison of the native mass of unmodified and modified BiP, whether from FICD-containing cells or from samples reacted with FICD and ATP in vitro, point to the addition of a single AMP moiety on any given molecule of BiP (Preissler et al., 2015b). Liquid chromatography coupled to mass spectrometry (LC-MS) of peptides arising from a digest with Arg-C revealed the presence of a single mono-AMPylated BiP^511-532^ peptide (m/z = 1375.15) (Preissler et al., 2015b) that was absent from samples of BiP that had never been AMPylated [(Preissler et al., 2015b) and Figure 3A, upper panel]. Crucially, this signal was abolished by exposure to wildtype FICD but not by exposure to the FICD^H363A^ mutant (Figure 3A, middle and lower panels). The loss of the AMPylated BiP^511-532^ peptide caused by FICD was matched by gain in peptides with the mass of the unmodified BiP^511-532^ (m/z = 1209.62) (Figure 3B) pointing to the ability of FICD to revert BiPT^518-AMP^ to its pre-modified state.

**Figure 3.**
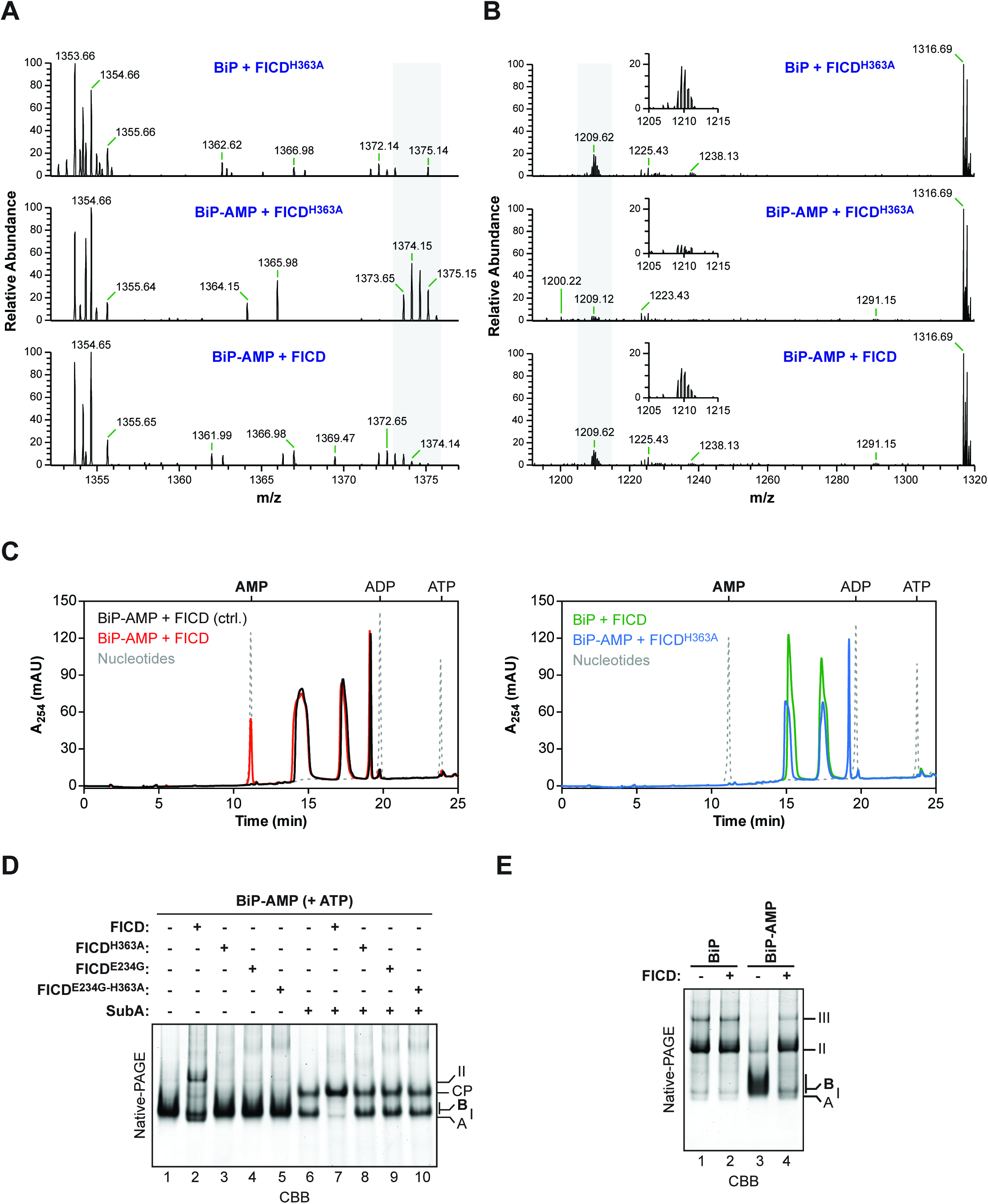
FICD-mediated de-AMPylation releases AMP and restores BiP to its pre-AMPylation state. (**A**) LC-MS spectra of peptides from a digest with Arg-C of unmodified BiP protein (upper spectrum) or in vitro AMPylated BiP (BiP-AMP; lower two spectra) incubated with wildtype FICD or the FICD^H363A^ mutant protein for 3 hours at 30°C. The isotopic series of doubly charged ions derived from the AMPylated proteolytic BiP fragment (BiP^511-532^; 1374.15 m/z, z = 2) are highlighted in grey. (**B**) As in “A” but showing the spectra of the unmodified BiP^511-532^ peptides (1209.62 m/z, z = 2; isotopic ensemble highlighted in grey and magnified in the inset). (**C**) Absorbance traces at 254 nm (A_254_ in milli absorbance units) of ion pair chromatograms of post-proteinaceous supernatants from samples of unmodified or in vitro AMPylated BiP after incubation with wildtype FICD or FICD^H363A^ (2 hours at 30°C). The peaks in the elution profile corresponding to known standards (ATP, ADP, and AMP in grey dotted lines) are provided as a reference. In the control reaction (black trace on the left panel) the FICD enzyme was added to BiP-AMP followed immediately by deproteination. (**D**) CBB-stained native-PAGE gel of purified in vitro AMPylated BiP that had been subsequently exposed to the indicated FICD enzymes in presence of ATP. The samples in lanes 6-10 were further exposed to SubA protease for 10 minutes at 25°C. The migration of the non-AMPylated monomeric ‘A’ form (enriched in the presence of ATP), BiP dimers (II), the AMPylated ‘B’ form, and nucleotide binding domain SubA cleavage product (CP) are noted. (**E**) As in “D”, non-AMPylated and AMPylated BiP, resolved by native-PAGE but in the absence of ATP. Note the recovery of BiP oligomers (labeled II & III) in the AMPylated sample that had been exposed to FICD (lane 4).

To examine the fate of the modifying nucleotide, protein-free supernatants of in vitro de-AMPylation reactions were examined by ion pair chromatography. Exposure of BiP^T518-AMP^ to wildtype FICD led to the emergence of a 254 nm absorbance peak that overlapped with the AMP marker (Figure 3C, red trace) and had indistinguishable absorption spectra by three-dimensional analysis (Figure 3 supplement 1). The AMP peak was conspicuously absent from reactions set up with wildtype FICD and non-AMPylated BiP (green trace) or AMPylated BiP and the inactive FICD^H363A^ mutant (blue trace). These experiments point to a role of FICD as a phosphodiesterase that liberates AMP from AMPylated BiP.

AMPylated BiP is locked into a low-substrate affinity, interdomain-coupled state that is relatively resistant to cleavage by SubA (Preissler et al., 2015b), a highly specific protease that cuts the interdomain linker of BiP (Paton et al., 2006). Native-PAGE revealed that exposure to wildtype FICD imparted sensitivity to SubA on AMPylated BiP (Figure 3D). In the absence of ATP, BiP forms higher order oligomers, a process that is impeded by AMPylation (Preissler et al., 2015a). Exposure to wildtype FICD led to oligomerization of the largely monomeric AMPylated form of BiP (Figure 3E, compare lanes 3 & 4). The FICD-mediated conversion of modified BiP to self-binding unmodified BiP was also noticeable in native gels of samples in presence of ATP; but only BiP dimers were detectable (Figure 3D, compare lanes 1 & 2). Thus, FICD restored BiP to its pre-AMPylated functional state.

FICD-mediated BiP AMPylation is specific for the intact full-length BiP, as the enzyme fails to recognize T518, when the latter is presented in the context of the isolated substrate-binding domain of BiP (Preissler et al., 2015b). To examine the substrate specificity of FICD's phosphodiesterase activity, the ability of SubA to cleave AMPylated BiP quantitatively (by prolonged incubation) was exploited. Exposure to FICD led to the time-dependent disappearance of the fluorescent signal from intact BiP^T518-AMP-FAM^. However, FICD was unable to remove the fluorescent moiety from the isolated substrate-binding domain of BiP^T518-AMP-FAM^ (Figure 4A). Kinetic analysis of FICD's phosphodiesterase activity revealed a substrate K_*M*_ of 15.58 ± 3.27 µM and a k_*cat*_ of 9.89 ± 0.87 × 10^−3^ sec^−1^ (Figure 4B & C). These observations are consistent with a specific but relatively slow enzyme, sensitive to the lower end of physiological fluctuations predicted in the concentration of its substrate.

**Figure 4.**
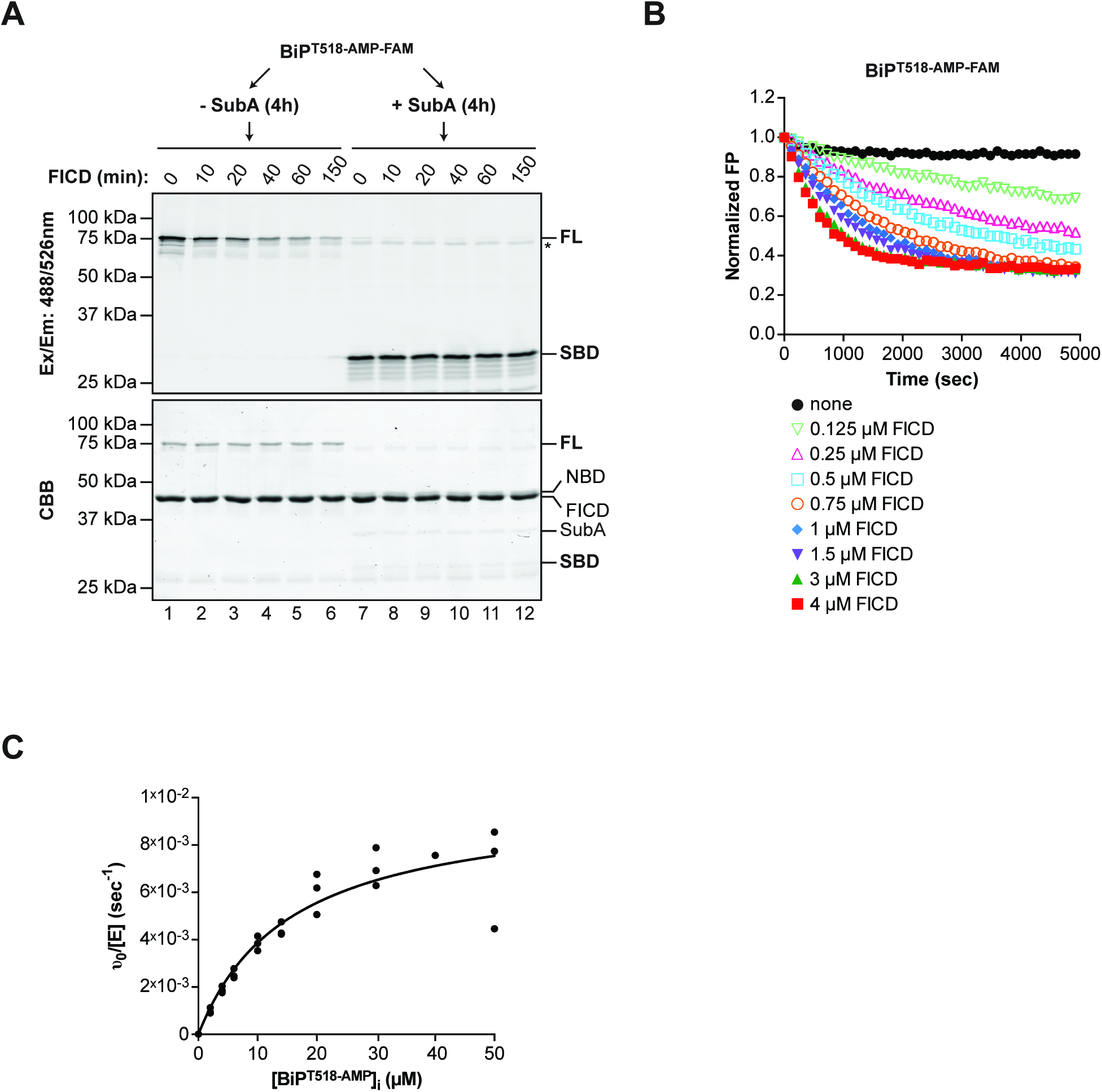
Enzymatic properties of the FICD de-AMPylase. (**A**) Fluorograph (excitation: 488 nm; emission 526 nm) and Coomassie stain of BiP AMPylated with FAM-labeled AMP (BiP^T518-AMP-FAM^) and exposed to wildtype FICD for the indicated time before SDS-PAGE. The BiP^AMP-FAM^ in samples loaded onto lanes 7-12 had been cleaved first with SubA protease, generating a substrate-binding domain fragment (SBD) labeled on BiP^T518^, before exposure to FICD. The location of the full-length BiP (FL) the isolated non-fluorescent nucleotide binding domain fragment (NBD), FICD and SubA are noted. The asterisk marks a high molecular weight fluorescent species distinct from fulllength BiP^AMP-FAM^ (FL) that is not a substrate for FICD-mediated de-AMPylation. Note the poor de-AMPylation of the SBD fragment. (**B**) Time-dependent plot of fluorescence polarization (FP) measurement of BiP^T518-AMP-FAM^ (80 nM) exposed to increasing concentrations of wildtype FICD (0.125 to 4 µM) as indicated. (**C**) Plot of the relation between initial substrate concentration [BiP^T518-AMP^]_i_ and the initial velocity of de-AMPylation derived from change in FP of BiP^T518-AMP-FAM^ (introduced as a tracer at an initial concentration of 17 nM). Wildtype FICD, [E] = 0.75 µM. Data points from three independent experiments (*n* = 3) and the nonlinear fit curve are shown (K_*M*_ = 15.58 µM ± SD 3.27 µM and k_*cat*_ = 9.89 × 10^−3^ sec^−1^ ± SD 0.87 × 10^−3^ sec^−1^).

The FICD-mediated BiP AMPylation/de-AMPylation cycle converts the cosubstrate ATP to AMP and pyrophosphate. Therefore, the production of AMP in reactions assembled with BiP, ATP, and combinations of FICD enzymes was used to evaluate the relative contribution of AMPylation and de-AMPylation to the apparent inability of wildtype FICD to promote a pool of AMPylated BiP (Figure 1 & 2C). No AMP was observed in reactions with the de-AMPylation defective FICD^E234G^, consistent with stability of BiP^T518-AMP^ (Figure 5A, blue trace), however, substantial amounts of AMP were produced over time when wildtype FICD was included alongside the hyperactive mutant FICD^E234G^ (Figure 5A, red trace and Figure 5B). Importantly, AMP was not produced in reactions with wildtype FICD alone (Figure 5A, black trace). These observations indicate that in the absence of other yet-to-be-identified cellular components the wildtype enzyme is locked in an AMPylation incompetent state, as suggested previously (Bunney et al., 2014; Engel et al., 2012). However, the wildtype enzyme freely de-AMPylates BiP and a single FICD^E234G^ mutation flips the activity of FICD from a constitutive de-AMPylase to a pure AMPylase.

**Figure 5.**
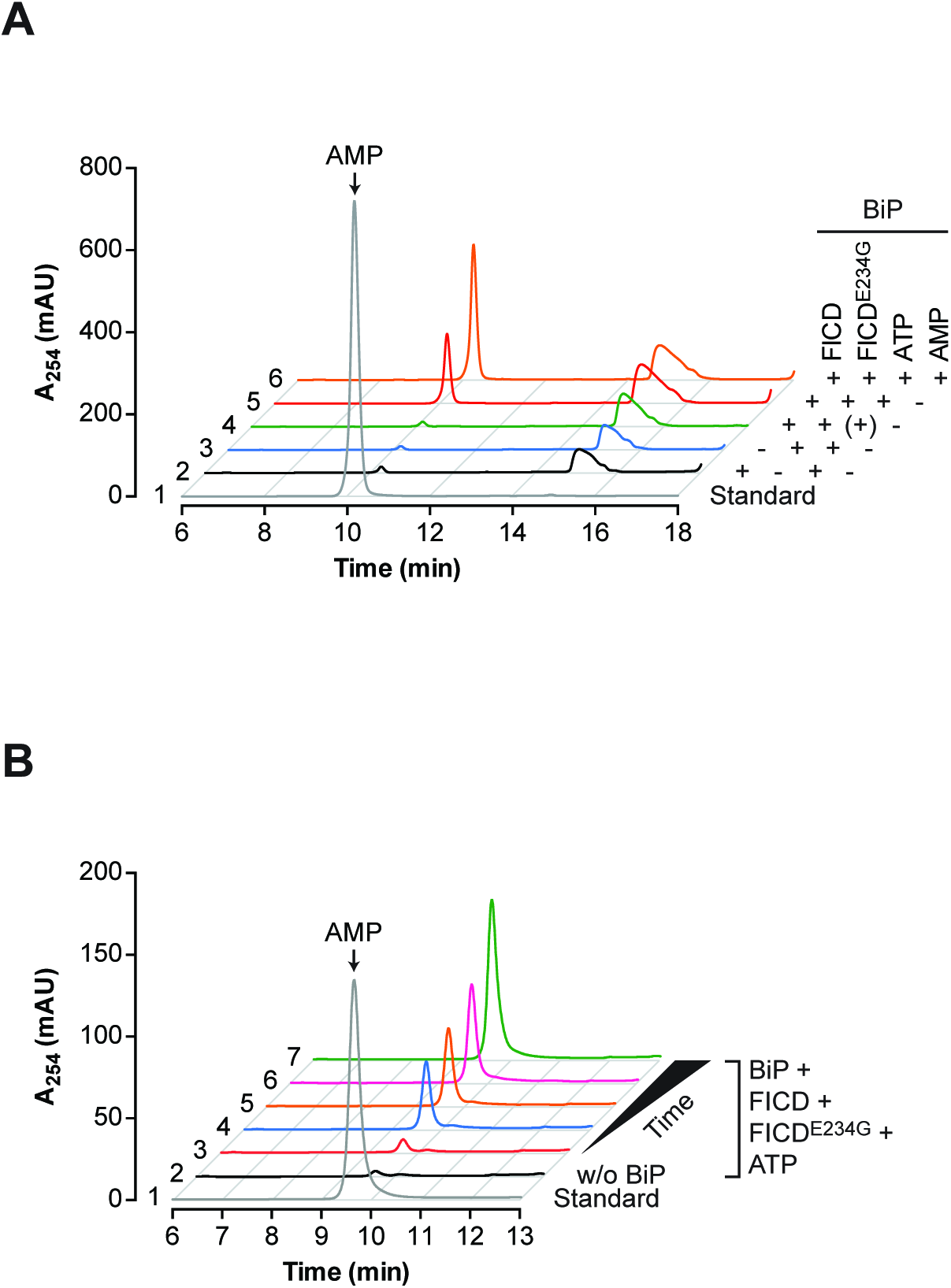
Engagement of E234 in the active site switches FICD from AMPylation to de-AMPylation. (**A**) Absorbance traces of ion pair chromatograms of post-proteinaceous supernatants from samples derived from the indicated enzymatic reactions (all reactions contained 20 µM BiP and 2 mM ATP and were incubated with or without 2 µM FICD and/or 2 µM FICD^E234G^ for 3 hours at 30°C). Shown is the AMP-containing region of the chromatogram. In the control reaction (trace 4, green) ATP was added followed immediately by deproteination. Trace 5 and 6 are of identical samples with the addition of AMP to trace 6 before chromatography (orange). Note that production of AMP is restricted to the reaction containing both FICD^E234G^ and wildtype FICD. (**B**) Absorbance traces (as above) of post-proteinaceous supernatants from samples derived from the indicated enzymatic reactions (20 µM BiP, 2 µM FICD, and 2 µM FICD^E234G^) incubated for 3 hours at 30°C. ATP (2 mM) was added at different times before deproteination (trace 3 = 0 min, trace 4 = 15 min, trace 5 = 30 min, trace 6 = 1 hour, traces 2 and 7 = 3 hours) to start the reactions. Shown is the AMP-containing region of the chromatogram. Trace 1 (grey line) represents the elution profile of an AMP standard. As a control, a reaction without BiP was included (trace 2, black line). Note the time-dependence of AMP production in reactions (3 to 7) containing FICD^E234G^, wildtype FICD and substrate BiP.

To explore the role of FICD as a BiP de-AMPylating enzyme in vivo, the effect of over-expression of the wildtype enzyme on the level of AMPylated BiP was measured in wildtype CHO-K1 cells. Exposure to cycloheximide rapidly led to the emergence of a strong signal of AMPylated BiP, detected by its characteristic mobility on native-PAGE gels (‘B’ form) (Preissler et al., 2015b). This signal was progressively attenuated by overexpression of wildtype FICD (Figure 6A), consistent with de-AMPylation of endogenous BiP by overexpressed FICD.

**Figure 6.**
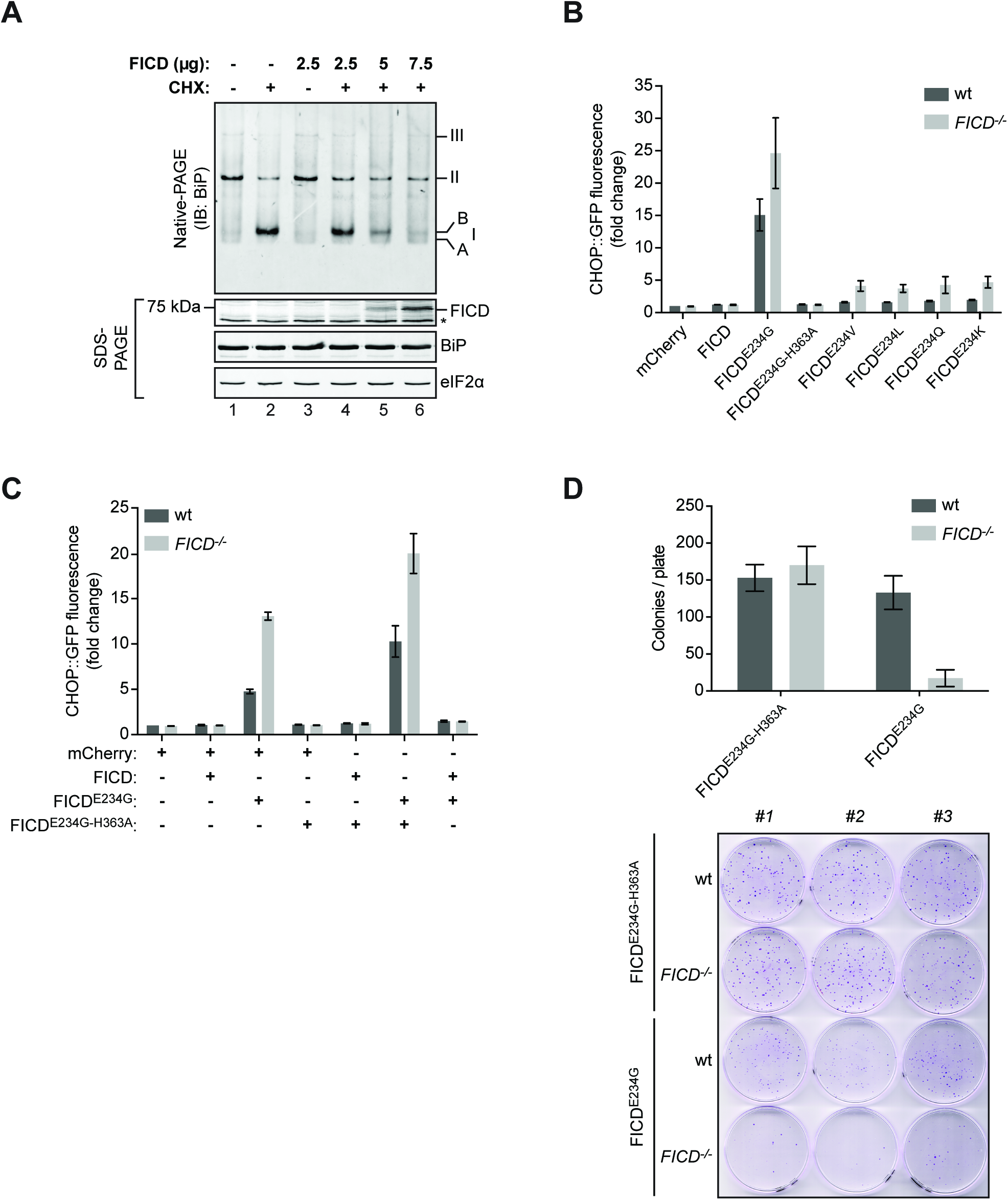
FICD counteracts BiP AMPylation in cells. (**A**) Immunoblot of endogenous BiP from CHO-K1 cells resolved by native-PAGE. The cells were transfected with the indicated amount of plasmid DNA encoding wildtype FICD and exposed to cycloheximide (CHX; 100 µg/mL, 3 hours) to promote AMPylated BiP. The AMPylated ‘B’ form of BiP is indicated (as are the other major species, see Figure 1 legend). Immunoblots of the same samples resolved by SDS-PAGE report on FICD, total BiP and total eIF2α (which also serves as a loading control). Data representative of three independent experiments are shown (*n* = 3). (**B**) Activity of an integrated *CHOP::GFP* UPR reporter in isogenic wildtype and FICD-deficient CHO-K1 cells following transfection with plasmids encoding the indicated FICD derivatives (marked by an mCherry transgene). Shown are the median values ± SD of the GFP fluorescent signal of the mCherry-positive cells from three independent experiments (fold change relative to wildtype cells transfected with control plasmid DNA encoding mCherry only; *n* = 3). (**C**) As in “B”, but cells were co-transfected with the indicated pairs of plasmids. Data from three independent experiments are shown (*n* = 3). (**D**) Bar diagram comparing outgrowth of puromycin-resistant colonies of wildtype and FICD-deficient CHO-K1 cells transduced with retrovirus (with a puromycin resistance maker) expressing a catalytically inactive FICD^E234G-H363A^ or de-AMPylation defective/AMPylation active FICD^E234G^. Shown are the mean values ± SD of the number of colonies per plate from three replicates (*n* = 3). Below is a photograph of the crystal violet stained colonies from the samples described above.

BiP potently represses UPR signaling (Bertolotti et al., 2000; Pincus et al., 2010), whereas inactivation of BiP, by enforced AMPylation, induces the UPR (Preissler et al., 2015b). A role for endogenous FICD in BiP de-AMPylation predicts more UPR activity in *FICD*^−/−^ cells targeted with de-AMPylation defective, AMPylation competent FICD derivatives than in similarly-targeted wildtype cells. To test this prediction, the intensity of the UPR was compared between isogenic wildtype and *FICD*^−/−^UPR reporter-bearing CHO-K1 cells transfected with plasmids encoding FICD derivatives. Expression of the catalytically-inactive FICD^E234G-H363A^ had no effect on UPR signaling, whereas de-AMPylation defective, AMPylation competent FICD derivatives with mutations at E234, reproducibly induced more UPR signaling in the *FICD*^−/−^ cells (Figure 6B & Figure 6 supplement 1). Conversely, co-expression of the de-AMPylation competent wildtype FICD markedly attenuated UPR signaling induced by the hyperactive FICD^E234G^ (Figure 6C & Figure 6 supplement 2). Treatment with the ER stress-inducing compounds tunicamycin or thapsigargin, or transfection of effector plasmids expressing the Cas9 nuclease and single guide RNAs that target the BiP gene promoted similar levels of UPR signaling in wildtype and *FICD*^−/−^cells (Figure 6 supplement 3), attesting to the selective sensitization by the *FICD*^−/−^ genotype towards effectors that inactivate BiP by AMPylation.

BiP inactivation has a large fitness cost (Paton et al., 2006; Preissler et al., 2015b). In keeping with a role for endogenous FICD in reversing BiP inactivation by AMPylation, wildtype CHO-K1 cells tolerated stable expression of a retrovirus encoding the de-AMPylation defective, AMPylation-competent FICD^E234G^ better than *FICD*^−/−^ mutant cells (Figures 6D). Together, these observations point to a role for endogenous FICD in reversing BiP AMPylation and restoring chaperone activity in cells.

## Discussion

FICD is both necessary and sufficient for BiP inactivation by AMPylation. The same enzyme is implicated here in removing this modification and in BiP reactivation. The side chain of a single residue, E234 determines which of the two opposing activities FICD will manifest in vitro. E234 of wildtype FICD blocks all AMPylation and renders the enzyme a pure, highly specific BiP de-AMPylase. Substitutions at E234, abolish the de-AMPylase activity of FICD and unmask, to varying degrees, its AMPylating activity.

FICD’s E234 lies at the tip of a regulatory helix conserved in other FIC enzymes (Garcia-Pino et al., 2014; Harms et al., 2016; Roy and Cherfils, 2015) (Figure 7). Engagement of the E234 side-chain in the active site sterically and electronically, delocalizes the terminal phosphates of the bound ATP to repress AMPylation (Bunney et al., 2014; Engel et al., 2012), explaining why, in absence of other factors, wildtype FICD is inactive as an AMPylase. However, the E234 side-chain is flexible, and while engaged in the active site can either form a salt bridge with FICD's R374 or retain its charged group free to engage in alternative reactions (PDB 4uO4 and 4uOU) such as BiP de-AMPylation.

**Figure 7.**
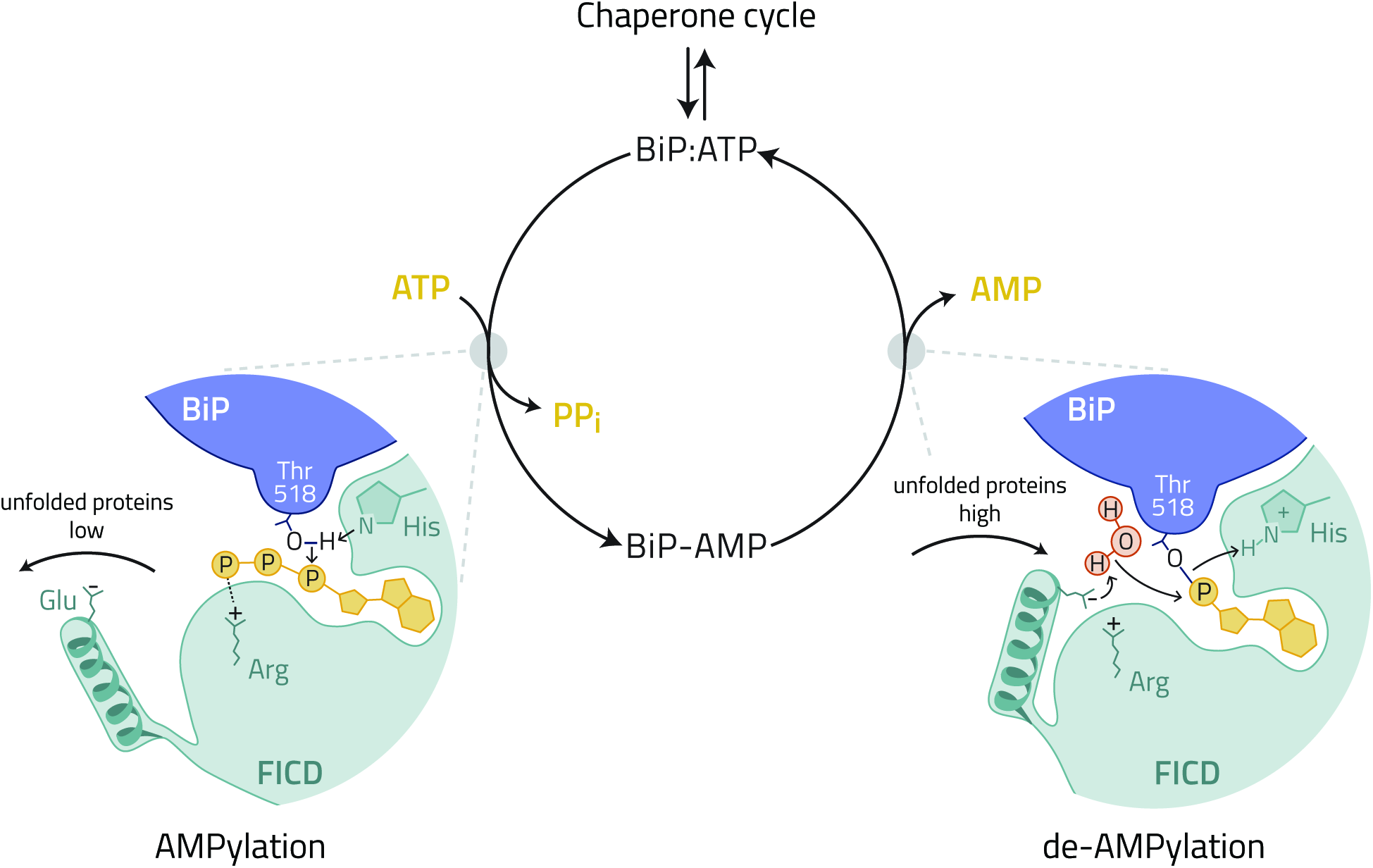
A hypothetical model depicting regulation of the FICD-mediated BiP AMPylation/de-AMPylation cycle by the disposition of E234. A cellular mechanism responsive to the changes in unfolded protein load is hypothesized to switch in FICD function by altering the position of E234. When unfolded proteins are scarce, active disengagement of FICD's E234 promotes the AMPylation-competent alignment of T518 of BiP, ATP and key residues from the active site (H363 and R374) (Bunney et al., 2014; Engel et al., 2012), inactivating BiP. When unfolded proteins are abundant FICD returns to its default state. The engaged E234 of FICD coordinates a water molecule to attack the phosphodiester bond between AMP and T518 of BiP. The essential role of FICD's H363 in de-AMPylation is plausibly attributed to protonation of the BiP T518 leaving group. De-AMPylation returns BiP to the chaperone cycle.

In vitro, the two forms of FICD, wildtype and E234G, represent the extremes of two opposing enzymatic activities. It is likely that other components, present in the ER, specify which of the two activities will prevail at any time, thus sparing the cell fruitless cycles of ATP-consuming BiP AMPylation and de-AMPylation. It is tempting to speculate that a mechanism exists for coupling the disposition of the E234-containing helix to the burden of unfolded protein in the ER, such that when this burden is high the E234 side-chain is engaged in the active site to block AMPylation and favor de-AMPylation. This appears to be the default conformation of pure FICD, explaining its inability to AMPylate BiP and its constitutive de-AMPylating activity in vitro. When the burden of unfolded proteins is low, a conformational switch in FICD disengages the E234 side chain to promote AMPylation. The mechanism behind this in vivo switch remains to be worked out, nonetheless the dominance of de-AMPylation over AMPylation when the wildtype enzyme is overexpressed, suggests that in the ER too the default state of E234 is to engage the active site and that an active mechanism, triggered in vivo when the burden of unfolded proteins is low, pries E234 from the active site converting wildtype FICD to an AMPylase. The machinery for switching FICD from de-AMPylation to AMPylation is absent from the in vitro assays conducted here but the consequences of its action are mimicked by the E234G mutation, which locks FICD in a constitutively AMPylating mode (Figure 7).The FIC domain is highly flexible in terms of substrate utilization. The bacterial FIC protein DOC has been observed both to phosphorylate threonine 382 of bacterial EF-Tu and to dephosphorylate the same residue. Analysis of the reaction products indicates that DOC mediated de-phosphorylation is achieved by thermodynamically-unfavored reversal of the phosphorylation reaction, regenerating a nucleotide tri-phosphate (Castro-Roa et al., 2013). By contrast the reaction products of FICD-mediated BiP de-AMPylation (unmodified BiP and free AMP) argue against simple mass action-driven enzymatic micro-reversibility and suggest instead that the active site of FICD is exploited for two interdependent, physiologically-antagonistic, chemically-distinct reactions.

AMPylation by FIC-domain enzymes employs a conserved HPFx(D/E)GN(G/K)R catalytic loop to position the attacking nucleophile in close proximity to the α-phosphate of the bound ATP substrate (Khater and Mohanty, 2015; Luong et al., 2010; Xiao et al., 2010). A co-crystal structure of the bacterial enzyme, IbpA, and its AMPylated target, Cdc42, indicates that the same pocket in the FIC active site can accommodate adenosine-phosphate when it is part of the ATP substrate and as part of the AMPylated Cdc42 product (Xiao et al., 2010). Both AMPylation and de-AMPylation by FICD require the conserved H363. Thus, a shared active site residue contributes to two antagonistic reactions initiated by different attacking nucleophiles.

In AMPylation, FICD mediates a concerted deprotonation and attack of the T518 hydroxyl of BiP on the α-phosphate of the bound ATP substrate (Bunney et al., 2014; Xiao et al., 2010). In de-AMPylation the hydroxyl of a water molecule, likely activated by E234, may attack the phosphodiester bond of the bound AMP. It is tempting to speculate that the essential role of FICD's H363 in both reactions reflects deprotonation of the attacking nucleophile in the AMPylation reaction and protonation of the BiP T518 leaving group in the de-AMPylation reaction.

In summary, the findings presented here are consistent with a model whereby BiP AMPylation evolved as a cellular buffer to fluctuations in unfolded ER proteins by acquisition of a single dual-functioning enzyme whose activity is switched in vivo by positioning a conserved regulatory residue. Because it responds directly to changes in unfolded protein load, without need for gene expression or protein synthesis, the machinery for executing this switch will teach us something about the most proximal steps in protein folding homeostasis in the ER.

## Materials and methods

### Plasmid construction

Supplementary file 1 lists the plasmids used in this study. Standard PCR and molecular cloning methods were used to generate DNA constructs and point mutations were introduced by PCR-based site-directed mutagenesis.

### Protein purification

N-terminally hexahistidine- (His6-) tagged wildtype and mutant Chinese hamster BiP proteins were expressed in M15 *E. coli* cells (Qiagen). The bacterial cultures were grown at 37°C to an optical density (OD_600 nm_) of 0.8 in LB medium supplemented with 50 µg/ml kanamycin and 100 µg/ml ampicillin and expression of recombinant protein was induced by the addition of 1 mM isopropylthio β-D-1-galactopyranoside (IPTG). The cells were further incubated at 37°C for 6 hours, harvested by centrifugation, and lysed with a high-pressure homogenizer (EmulsiFlex-C3, Avestin) in buffer A [50 mM Tris-HCl pH 7.5, 500 mM NaCl, 1 mM MgCl_2_, 0.2% (v/v) Triton X-100, 10% (v/v) glycerol, 20 mM imidazole] containing protease inhibitors [2 mM phenylmethylsulphonyl fluoride (PMSF), 4 µg/ml pepstatin, 4 µg/ml leupeptin, 8 µg/ml aprotinin] and 0.1 mg/ml DNaseI. The obtained lysates were cleared by centrifugation for 30 minutes at 25,000 *g* and incubated with 1 ml Ni-NTA agarose (Qiagen) per 1 l of expression culture for 2 hours at 4°C. The matrix was transferred to a column and washed five times with 20 bed volumes of buffer A containing 5 mM β-mercaptoethanol and supplemented sequentially with (i) 30 mM imidazole, (ii) 1% (v/v) Triton X-100, (iii) 1 M NaCl, (iv) 5 mM Mg^2+^-ATP, or (v) 0.5 M Tris-HCl pH 7.5. Bound BiP proteins were eluted in buffer B [50 mM HEPES-KOH pH 7.5, 300 mM NaCl, 10% (v/v) glycerol, 5 mM β-mercaptoethanol, 250 mM imidazole] and dialyzed against HKM buffer [50 mM HEPES-KOH pH 7.4, 150 mM KCl, 10 mM MgCl2]. The proteins were concentrated using centrifugal filters (Amicon Ultra, 30 kDa MWCO; Merck Millipore), snap-frozen in liquid nitrogen, and stored at -80°C.

Bacterial expression and purification of N-terminally GST-tagged wildtype and mutant FICD proteins was performed according to (Preissler et al., 2015b) with modifications. The FICD-encoding DNA constructs were transformed into C3013 BL21 T7 Express *lysY/I*^*q*^ *E. coli* cells (New England BioLabs) and cultures of single clones were grown at 37°C in LB medium containing 100 µg/ml ampicillin. At an OD_600 nm_ of 0.8 the cultures were shifted to 20°C and expression was induced with 0.5 mM IPTG. After incubation for 16 hours the cells were harvested and lysed as described above in lysis buffer [50 mM Tris-HCl pH 7.5, 500 mM NaCl, 1 mM MgCl_2_, 2 mM dithiothreitol (DTT), 0.2% (v/v) Triton X-100, 10% (v/v) glycerol] containing protease inhibitors and DNaseI. The lysates were cleared by centrifugation for 30 minutes at 25,000 *g* and incubated with 0.7 ml Glutathione Sepharose 4B (GE Healthcare) per 1 l of expression culture for 2 hours at 4°C. The beads were washed with 20 ml wash buffer C [50 mM Tris-HCl pH 7.5, 500 mM NaCl, 1 mM DTT, 0.2% (v/v) Triton X-100, 10% (v/v) glycerol] containing protease inhibitors, 20 ml wash buffer D [50 mM Tris-HCl pH 7.5, 300 mM NaCl, 10 mM MgCl_2_, 1 mM DTT, 0.1% (v/v) Triton X-100, 10% (v/v) glycerol] containing protease inhibitors and 20 ml wash buffer D sequentially supplemented with (i) 1% (v/v) Triton X-100, (ii) 1 M NaCl, (iii) 3 mM ATP, or (iv) 0.5 M Tris-HCl pH 7.5. Bound proteins were eluted in elution buffer [50 mM HEPES-KOH pH 7.4, 100 mM KCl, 4 mM MgCl_2_, 1 mM CaCl_2_, 0.1% (v/v) Triton X-100, 10% (v/v) glycerol, 40 mM reduced glutathione] and concentrated protein solutions were frozen in liquid nitrogen and stored at -80°C. For preparation of wildtype FICD without the GST-tag the protein was eluted from glutathione-Sepharose in elution buffer without Triton X-100 and with 1.5 mM DTT. TEV protease was added in a 100:1 (protein-to-TEV) molar ratio and after incubation for 16 hours at 4°C the proteins were passed over a size-exclusion chromatography column (Superdex 200 10/300 GL; GE Healthcare) in HKMG buffer [50 mM HEPES-KOH pH 7.4, 150 mM KCl, 10 mM MgCl_2_, 10% glycerol]. The FICD protein-containing fractions were pooled, concentrated, and frozen in aliquots.

### Purification of in vitro AMPylated BiP proteins

AMPylated BiP proteins were prepared as described previously (Preissler et al., 2015b), with modifications. In brief, 20 mg of purified wildtype BiP or substrate-binding deficient BiP^V461F^ mutant protein (Petrova et al., 2008) was in vitro AMPylated for 4 hours at 30°C with 0.25 mg bacterially expressed FICD^E234G^ in presence of 3 mM ATP in buffer E [25 mM HEPES-KOH pH 7.4, 100 mM KCl, 10 mM MgCl_2_, 1 mM CaCl_2_, 0.1% (v/v) Triton X-100]. Afterwards, BiP proteins were bound to 400 µl Ni-NTA agarose affinity matrix for 30 minutes at 25°C, washed with buffer E, and eluted in buffer E containing 350 mM imidazole for 45 minutes at 25°C. The eluate was concentrated and passed over a Centri•Pure P25 desalting column (emp BIOTECH) equilibrated in HKMG buffer. The protein-containing fractions were pooled, concentrated, frozen in liquid nitrogen, and stored at -80°C. BiP was quantitatively AMPylated as judged by the conversion of BiP oligomers into the modified monomeric ‘B’ form on a native-PAGE gel. The modified BiP proteins were used for in vitro de-AMPylation assays and mass spectrometry analysis (see below). Unmodified BiP prepared from a parallel mock AMPylation reaction without enzyme served as a control.

### In vitro AMPylation and de-AMPylation assays

Unless stated otherwise in vitro AMPylation and de-AMPylation reactions were performed in AMPylation buffer [25 mM HEPES-KOH pH 7.4, 100 mM KCl, 4 mM MgCl_2_, 1 mM CaCl_2_, 0.1% (v/v) Triton X-100].

Radioactive in vitro AMPylation (Figure 2C) reactions were set up in a final volume of 37.5 µl containing 1 µM of ATP hydrolysis-deficient mutant BiP protein (BiP^T229A^) (Gaut and Hendershot, 1993), 0.1 µM wildtype or mutant FICD proteins, 40 µM ATP, and 0.023 MBq α-^32^P-ATP (EasyTide; Perkin Elmer). The reactions were started by addition of the nucleotides and incubated at 25°C. After 3 and 10 minutes of incubation 15 µl were removed from each reaction, respectively, supplemented with 5 µl SDS sample buffer, heated for 5 minutes at 75°C and loaded on a SDSPAGE gel. Gels were stained with Coomassie and the radioactive signals were detected with a Typhoon Trio imager (GE Healthcare) upon overnight exposure of the dried gels to a storage phosphor screen.

For radioactive de-AMPylation experiments (Figure 2A) BiP was first AMPylated in vitro with α-^32^P-ATP and then re-purified. Therefore, 6 µg of purified BiP was preincubated with 15 µM ATP in a final volume of 20 µl in AMPylation buffer for one minute at 25°C before 1.85 MBq α-^32^P-ATP and 2 µg FICD^E234G^ was added. The mixture was incubated for 10 minutes at 25°C and another 12 µg of BiP was added. After further incubation for 50 minutes the reaction was diluted with 200 µl of highsalt AMPylation buffer [25 mM HEPES-KOH pH 7.4, 500 mM KCl, 4 mM MgCl_2_, 1 mM CaCl_2_, 0.1% (v/v) Triton X-100] and 1 mM ATP was added. BiP was then bound to 20 µl Ni-NTA agarose beads for 30 minutes at 20°C. The beads were washed with 500 µl high-salt AMPylation buffer containing 1 mM ATP and three times with highsalt AMPylation buffer. Bound proteins were eluted in 100 µl AMPylation buffer containing 400 mM imidazole (pH 7.4) for 30 minutes at 20°C. The eluate was split in two fractions of 50 µl and each fraction was passed through a Sephadex G-50 MicroSpin column (illustra AutoSeq G-50; GE Healthcare) equilibrated with AMPylation buffer, and the recovered proteins were frozen in aliquots until the de-AMPylation experiment. The de-AMPylation reactions were carried out at 23°C in a final volume of 15 µl in AMPylation buffer containing 1.3 µg non-radioactive AMPylated wildtype BiP supplemented with trace amounts of radiolabelled AMPylated BiP (≈ 0.1 µg). The reactions were started at different time points by addition of 0.13 µg wildtype or mutant FICD. At the end of the experiment 5 µl SDS sample buffer were added to each reaction, proteins were denatured for 5 minutes at 75°C, and 15 µl of each sample were applied to SDS-PAGE. After separation, the proteins were stained with Coomassie to confirm equal loading and radioactive signals were detected by autoradiography as described above.

The non-radioactive de-AMPylation reactions shown in Figure 3D contained 1 µg/μl purified AMPylated wildtype BiP, 0.1 µg/μl wildtype or mutant FICD, and were incubated for 90 minutes at 30°C in a final volume of 25 µl in presence of 3 mM ATP. Afterwards, each reaction was divided in two samples, one of which was treated for 10 minutes with 0.06 µg/μl SubA protease at 25°C, whereas the other remained untreated. The samples were then supplemented with native sample buffer and analysed immediately by native-PAGE and Coomassie staining (see below). The de-AMPylation reactions shown in Figure 3E were performed in HKM buffer and contained 1 µg/μl purified AMPylated or unmodified wildtype BiP and were incubated without or with 0.05 µg/μl wildtype FICD for 30 minutes at 30°C in absence of ATP (to allow re-formation of BiP oligomers) before analysis by native-PAGE.

The reactions shown in Figure 2B contained 3 µg/μl protein from lysates of untreated wildtype CHO-K1 cells or cells treated for 3 hours with CHX, and 0.15 µg/μl purified wildtype or mutant FICD (see above). The reactions were started by addition of purified FICD and incubated at 30°C. After 15 minutes, the reactions were diluted 1:10 in IEF lysis buffer and analysed by IEF (see below).

### Mass spectrometry

For mass spectrometry analysis purified in vitro AMPylated or unmodified wildtype BiP at 16 µM was incubated with 0.8 µM wildtype FICD or catalytically inactive mutant FICD^H363A^ for 3 hours at 30°C. Afterwards, the proteins were denatured with SDS sample buffer, heated for 5 minutes at 75°C, and separated by SDS-PAGE. The gels were stained with Coomassie, destained, and the bands at 75 kDa corresponding to BiP protein were excised. The proteins were then reduced, alkylated, and digested “in-gel” with Arg-C protease. The obtained peptides were analyzed by LC-MS/MS using a Q Exactive mass spectrometer (Thermo Fischer) coupled to a RSLC 3000 UHPLC. The data were processed with Proteome Discoverer 1.4 using Sequest to search a Uniprot E. coli database (downloaded 29/04/15, 4377 entries) with the sequence of Chinese hamster (*Cricetulus griseus*) BiP added. Oxidation (M) and AMPylation (S/T) were set as variable modifications and carbamidomethylation (C) as a fixed modification. FDR calculations were performed by Percolator and peptides were filtered to 1%.

### Ion pair chromatography (IPC)

To detect the leaving group of the FICD-mediated de-AMPylation reaction, reversed-phase IPC was performed (Figure 3C). Purified in vitro AMPylated or unmodified BiP proteins at 65 µM were exposed to wildtype FICD or mutant FICD^H363A^ proteins at 6.5 µM in HKM buffer in a final volume of 30 µl for 2 hours at 30°C. At the end of the incubation time the reactions were stopped by addition of 10 µl of 4 M perchloric acid (PCA). As a negative control AMPylated BiP was incubated in parallel for 2 hours without enzyme and FICD was added directly before mixing with PCA. After incubation for 5 minutes on ice the samples were centrifuged at 21,000 *g* for 2 minutes at 4°C and 32 µl of the supernatants were mixed with 20 µl of 2 M potassium hydroxide (KOH). The pH of the samples was neutralized, the precipitates were sedimented by centrifugation for 15 minutes at 21,000 *g* at 4°C, and the cleared supernatant were equilibrated to room temperature before analysis by IPC. For that, 20 µl of each sample were injected onto a Poroshell 120 EC-C18 HPLC column (3 x 150 mm, 2.7 µm; Agilent Technologies) connected to a UHPLC Guard column (Agilent Technologies). Buffers A [H_2_O+10 mM tetrabutylammonium hydroxide (TBAH)+10 mM potassium dihydrogen phosphate (KH_2_PO_4_)] and B [methanol (CH_3_OH)+10 mM TBAH] were used as a mobile phase. The runs were performed at a constant flow rate of 0.4 ml/min at room temperature using the following gradient: 5% to 50% B in 25 minutes, hold for 2 minutes at 50% B, ramp to 95% B in 0.1 minute, hold for 7 minutes, 95% to 5% B in 1 minute, hold for 5 minutes at 5% B (reequlibration to basal). Nucleotide absorbance traces at 254 nm (A_254 nm_) were recorded and plotted against elution time. A nucleotide standard was applied in each experiment to determine the retention times of ATP, ADP and AMP.

Coupled AMPylation/de-AMPylation reactions (Figures 3D and S2) contained 20 µM ATP hydrolysis-deficient BiP^T229A^, 2 µM wildtype FICD, 2 µM FICD^E234G^, and 2 mM ATP in HKM buffer as indicated and were incubated for 3 hours at 30°C before deproteination and HPLC analysis as described above.

### Fluorescence polarization assay

The probe to measure the kinetics of FICD-mediated de-AMPylation by fluorescence polarization (FP) assays was generated as follows: Purified active FICD^E234G^ at 10 µM was pre-incubated with 2 mM 5-FAM-labeled N^6^-(6-Amino)hexyl-adenosine-5’-triphosphate (ATP-FAM; Jena Bioscience) in HKMG buffer for 5 minutes at 25°C before BiP^V461F^ protein at 10 µM was added to yield a final reaction volume of 50 µl. The substrate binding- and oligomerisation-deficient BiP^V461F^ mutant was used to avoid effects caused by BiP oligomerisation on fluorescence polarization measurements. The AMPylation reaction was allowed to proceed for 2 hours at 25°C. BiP protein was then bound to Ni-NTA agarose beads in presence of 0.01% (v/v) Triton X-100 and after several wash steps in the same buffer proteins were eluted in HKMG containing 250 mM imidazole. The eluted proteins were passed over a desalting column equilibrated in HKMG and aliquots were frozen in liquid nitrogen and stored at -80°C.

To confirm covalent modification of BiP^V461F^ with AMP-FAM the eluted proteins were incubated without or with 60 ng/μl SubA protease for an elongated period of time (4 hours) in AMPylation buffer at 30°C. The proteins were then denatured in SDS sample buffer for 5 minutes at 75°C and separated by SDS-PAGE. The fluorescence signals were detected with a Typhoon Trio imager (GE Healthcare; λ_ex_ = 488 nm, λ_em_ = 526 nm) before the gels were stained with Coomassie to visualize proteins (Figure S1A). To compare de-AMPylation of full-length BiP and the isolated substrate binding domain (Figure 4A) AMP-FAM modified BiP^V461F^ (BiP^V461F-AMPFAM^) was first cleaved for 4 hours with SubA and then exposed to 0.75 µM wildtype FICD protein for the indicated periods of time before denaturation and SDS-PAGE analysis as described above.

FP assays were performed in 384 well polysterene microplates (black, flat bottom, μCLAER; greiner bio-one) by monitoring fluorescence polarization of FAM (λ_ex_ = 485 nm, λ_em_ = 535 nm) using a TECAN plate reader (Infinite F500, *G*-factor≈0.82). The reactions were performed in a final sample volume of 30 µl and started by addition of either enzyme or substrate. To measure enzyme kinetics at excess of substrate the reactions contained the same concentration of enzyme (wildtype FICD protein without the GST-tag; [E] = 0.75 µM) and trace amounts of fluorescent BiP^V461F-AMP-FAM^ probe (17 nM) and were supplemented with increasing concentrations of non-fluorescent BiP^V461F-AMP^ ([S] = 2 µM to 50 µM), assuming that both were de-AMPylated with similar efficiency (Figure 4C). All reactions were performed at 30°C in AMPylation buffer. Substrate-only controls were included in each experiment for correction of the test sample values. The linear initial rates in millipolarization units per second (mP/sec) were converted to reaction velocity (μM/sec) using Equation 1, where ΔFP/Δt is the initial rate, ΔFP_c_ is the absolute change in FP signal at completion of the reaction, and [S] is the substrate concentration.

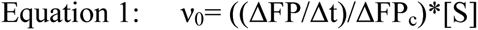

To calculate the K_m_ for FICD-mediated de-AMPylation of BiP^V461F-AMP^ the Michaelis-Menten equation (Michaelis et al., 2011) was fitted to a plot of the initial reaction velocities against the substrate concentrations from three independent experiments (*n* = 3). Data analysis was performed using Prism 6.0e (GraphPad Software, Inc.).

### Cell culture and mammalian cell lysates

All cells were grown on tissue culture dishes or multi-well plates (Corning) at 37°C and 5% CO_2_. CHO-K1 cells (ATCC CCL-61) were cultured in Nutrient mixture F-12 Ham (Sigma) supplemented with 10% (v/v) serum (FetalClone II; HyClone), 1 x Penicillin-Streptomycin (Sigma) and 2 mM L-glutamine (Sigma). The CHO-K1 *FICD*^−/−^ cell lines used in this study were described previously (Preissler et al., 2015b). HEK293T cells (ATCC CRL-3216) were cultured in Dulbecco's Modified Eagle's Medium (Sigma) supplemented as described above.

Experiments were performed at cell densities of 60-90% confluence. Cells were treated with drugs at the following final concentrations: 100 µg/ml cycloheximide (Sigma), 2.5 µg/ml tunicamycin (Melford), 0.5 µM thapsigargin (Calbiochem), and 6-8 µg/ml puromycin (Calbiochem). All drugs were first diluted in fresh, pre-warmed medium and then applied to the cells by medium exchange.

Cell lysis was performed as described in (Preissler et al., 2015a) with modifications. In brief, mammalian cells were cultured on 10 cm dishes and treated as indicated and/or transfected using Lipofectamine LTX with 5 µg plasmid DNA unless indicated otherwise, and allowed to grow for 24 hours. Before lysis the dishes were placed on ice, washed with ice-cold PBS, and cells were detached in PBS containing 1 mM EDTA using a cell scraper. The cells were sedimented for 5 minutes at 370 *g* at 4°C and lysed in HG lysis buffer [20 mM HEPES-KOH pH 7.4, 150 mM NaCl, 2 mM MgCl_2_, 10 mM D-glucose, 10% (v/v) glycerol, 1% (v/v) Triton X-100] containing protease inhibitors (2 mM PMSF, 4 µg/ml pepstatin, 4 µg/ml leupeptin, 8 µg/ml aprotinin) with 100 U/ml hexokinase (from *Saccharomyces cerevisiae* Type F-300; Sigma) for 10 minutes on ice. The lysates were cleared for 10 minutes at 21,000 *g* at 4°C. BIO-RAD protein assay reagent (BioRad) was used to determine the protein concentrations of lysates followed by normalization. For analysis by SDS-PAGE, SDS sample buffer was added to the lysates and proteins were denatured by heating for 10 minutes at 70°C before separation on 12.5% SDS polyacrylamide gels. To detect endogenous BiP by native-PAGE the lysate samples were loaded immediately on native gels (see below).

### Native polyacrylamide gel electrophoresis (native-PAGE)

Non-denaturing native-PAGE was performed as described (Preissler et al., 2015a). Briefly, Tris-glycine polyacrylamide gels (4.5% stacking gel and a 7.5% separation gel) were used to separate purified proteins or proteins from mammalian cell lysates to detect BiP oligomers. The separation was performed in running buffer (25 mM Tris, 192 mM glycine, pH ∼8.8) at 120 V for 2 hours. Afterwards, the proteins were visualized by staining with InstantBlue Coomassie solution (expedeon) or transferred to a polyvinylidene difluoride (PVDF) membrane in blotting buffer (48 mM Tris, 39 mM glycine; pH ∼9.2) supplemented with 0.04 (w/v) SDS for 16 hours at 30 V for immunodetection. The membrane was washed for 20 minutes in blotting buffer (without SDS) supplemented with 20% (v/v) methanol before blocking. Seven μg of purified BiP protein was loaded per lane to detect purified BiP proteins by Coomassie staining and volumes of lysates corresponding to 30 µg of total protein were loaded per lane to detect endogenous BiP from CHO-K1 cell lysates by immunoblotting.

### Immunoblot analysis

After separation by SDS-PAGE or native-PAGE (see above) the proteins were transferred onto PVDF membranes. The membranes were blocked with 5% (w/v) dried skimmed milk in TBS (25 mM Tris-HCl pH 7.5, 150 mM NaCl) and incubated with primary antibodies followed by IRDye fluorescently labeled secondary antibodies (LiCor). The membranes were scanned with an Odyssey near-infrared imager (LiCor). Primary antibodies and antisera against hamster BiP [chicken anti-BiP; (Avezov et al., 2013)], eIF2α [mouse anti-eIF2α; (Scorsone et al., 1987)], and FICD [chicken anti-FICD (Preissler et al., 2015b)] were used.

### Isoelectric focusing (IEF)

Analysis of lysates from mammalian cells by IEF was performed as described previously (Preissler et al., 2015b). Cells were grown in 10 cm dishes to approximately 90% confluence and treated with cycloheximide for 3 hours. Afterwards, the cells were washed with ice-cold TBS, resuspended in 1 ml TBS, sedimented by centrifugation, and lysed in 30 x its packed cell pellet volume of IEF lysis buffer [8.8 M urea, 5% (w/v) CHAPS, 1 µM sodium pyrophosphate, 2 mM imidodiphosphate, 50 mM DTT, 2% (v/v) Pharmalyte] at room temperature for 5 minutes. The lysates were centrifuged at 21,000 *g* for 10 minutes at room temperature and the resulting supernatants were centrifuged again for 60 minutes. The cleared lysates were passed over a Sephadex G-50 MicroSpin columns equilibrated with IEF sample buffer [8 M urea, 5% (w/v) CHAPS, 50 mM DTT, 2% (v/v) Pharmalyte (pH 4.5-5.4; GE Healthcare)], and 15 µl loaded on a 3.75% polyacrylamide gel containing 8.8 M urea, 1.25% (w/v) CHAPS, and 5% (v/v) Pharmalyte. The wells were overlaid with 0.5 M urea and 2% (v/v) Pharmalyte solution before the run. The anode buffer was 10 mM glutamic acid and the cathode buffer was 50 mM histidine. The run was performed as follows: 100 V for 10 minutes, 250 V for 1 hour, 300 V for 1 hour, 500 V for 30 minutes. The proteins were then transferred to a nitrocellulose membrane for 3 hours at 300 mA in blotting buffer [25 mM Tris-HCl pH 9.2, 190 mM glycine, 0.01% (w/v) SDS, 10% (v/v) methanol] and BiP was detected as described above.

### Flow cytometry

FICD (wildtype vs. mutants) overexpression-dependent induction of unfolded protein response signalling was analyzed by transient transfection of wildtype and *FICD*^−/−^ *CHOP::GFP* CHO-K1 UPR reporter cell lines with plasmid DNA encoding the FICD protein and mCherry as a transfection marker, using Lipofectamine LTX as described previously (Preissler et al., 2015b); 2 µg DNA were used in Figure 6B and 1 µg each plasmid were used in Figure 6C. Thirty-six hours after transfection cells were washed with PBS and collected in PBS containing 4 mM EDTA, and single cell fluorescent signals (20,000/sample) were analyzed by dual-channel flow cytometry with an LSRFortessa cell analyzer (BD Biosciences). GFP and mCherry fluorescence was detected with excitation laser 488 nm, filter 530/30, and excitation laser 561, filter 610/20, respectively. Data were processed using FlowJo and median reporter (in Q1 and Q2 in Figures 6B and 6C) analysis was performed using Prism.

The sensitivity to UPR induction in wildtype and FICD-deficient CHO-K1 CHOP::GFP UPR reporter cell lines was analyzed by treating cells with UPR-inducing compounds, tunicamycin or thapsigargin, for 16 hours before analysis, or transient transfection with 2 µg plasmid DNA encoding the Cas9 nuclease and single guide RNAs targeting hamster BiP. The two single guide RNA sequences (plasmids UK1857 and UK1858; Supplementary file 1) for targeting the second exon of *Cricetulus griseus* (Chinese hamster) BiP (*HSPA5*) were selected from the CRISPy database [URL: http://staff.biosustain.dtu.dk/laeb/crispy/, (Ronda et al., 2014)] and duplex DNA oligonucleotides of the sequences were inserted into the pSpCas9(BB)-2A-mCherry_V2 plasmid (plasmid UK1610; Supplementary file 1) as described (Ran et al., 2013). Four days after transfection cells were analysed by flow cytometry as described above.

### Production of VSV-G retroviral virus in HEK293T cells and infection of CHO-K1 cells

The effect of expression of de-AMPylation-defective/AMPylation-active FICD^E234G^ on mammalian cell fitness was analyzed by colony outgrowth of wildtype and *FICD*^−/−^CHO-K1 cell lines targeted with puromycin-resistant retrovirus expressing FICD^E234G^. For that HEK293T cells were split onto 6 cm dishes 24 hours prior to co-transfection of pBABE Puro plasmids (Morgenstern and Land, 1990) encoding FICD^E234G^ or inactive FICD^E234G-H363A^ (plasmids UK1852 and UK1853, respectively; Supplementary file 1) with VSV-G retroviral packaging vectors, using TransIT-293 Transfection Reagent (Mirus) according to the manufacturer's instructions. Sixteen hours after transfection, medium was changed to medium supplemented with 1% BSA (Sigma). Retroviral infections were performed following a 24 hours incubation by diluting 0.45 µm filter-sterilized cell culture supernatants at a 1:2 ratio into CHOK1 cells medium supplemented with 10 µg/ml polybrene (4 ml final volume) and adding this preparation to wildtype or *FICD*^−/−^ CHO-K1 cells (3 x 10^5^ cells seeded onto 6 cm dishes 24 hour prior to infection). Infections proceeded for 8 hours, after which viral supernatant was replaced with fresh medium. Forty-eight hours later cells were split into medium supplemented with 6 µg/ml puromycin, and 24 hours afterwards this was changed to medium supplemented with 8 µg/ml puromycin. The medium was changed every third day until puromycin-resistant colonies were visible. Colonies remaining on the dishes 9 days after infection were visualized by crystal violet stain and counted.

## Author Contributions

**Steffen Preissler:** Conceived, designed and led the project. Conducted in vitro experiments, analysis and interpretation of data, drafting and revising the article.

**Claudia Rato:** Designed, conducted and interpreted the in vivo experiments, and contributed to drafting and revising the article.

**Luke Perera:** Contributed to protein purification and fluorescence polarization experiments and revised the manuscript.

**Vladimir Saudek:** Provided valuable insights and discussions and revised the manuscript.

**David Ron:** Oversaw the project, conception and design, construction of plasmid DNA, analysis and interpretation of data, drafting and revising the article.

## Acknowledgements

We thank R. Antrobus from CIMR mass spectrometry, R. Schulte and the CIMR flow cytometry team for assistance, Heather P. Harding, Niko Amin-Wetzel, and Joseph Chambers (CIMR) for advice and comments on the manuscript and Claudia Flandoli (Cambridge, UK) for the cartoon. Supported by grants from the Wellcome Trust (Wellcome 200848/Z/16/Z and a strategic award Wellcome 100140). D.R. is a Wellcome Trust Principal Research Fellow.

---

**Figure 2 supplement 1.**
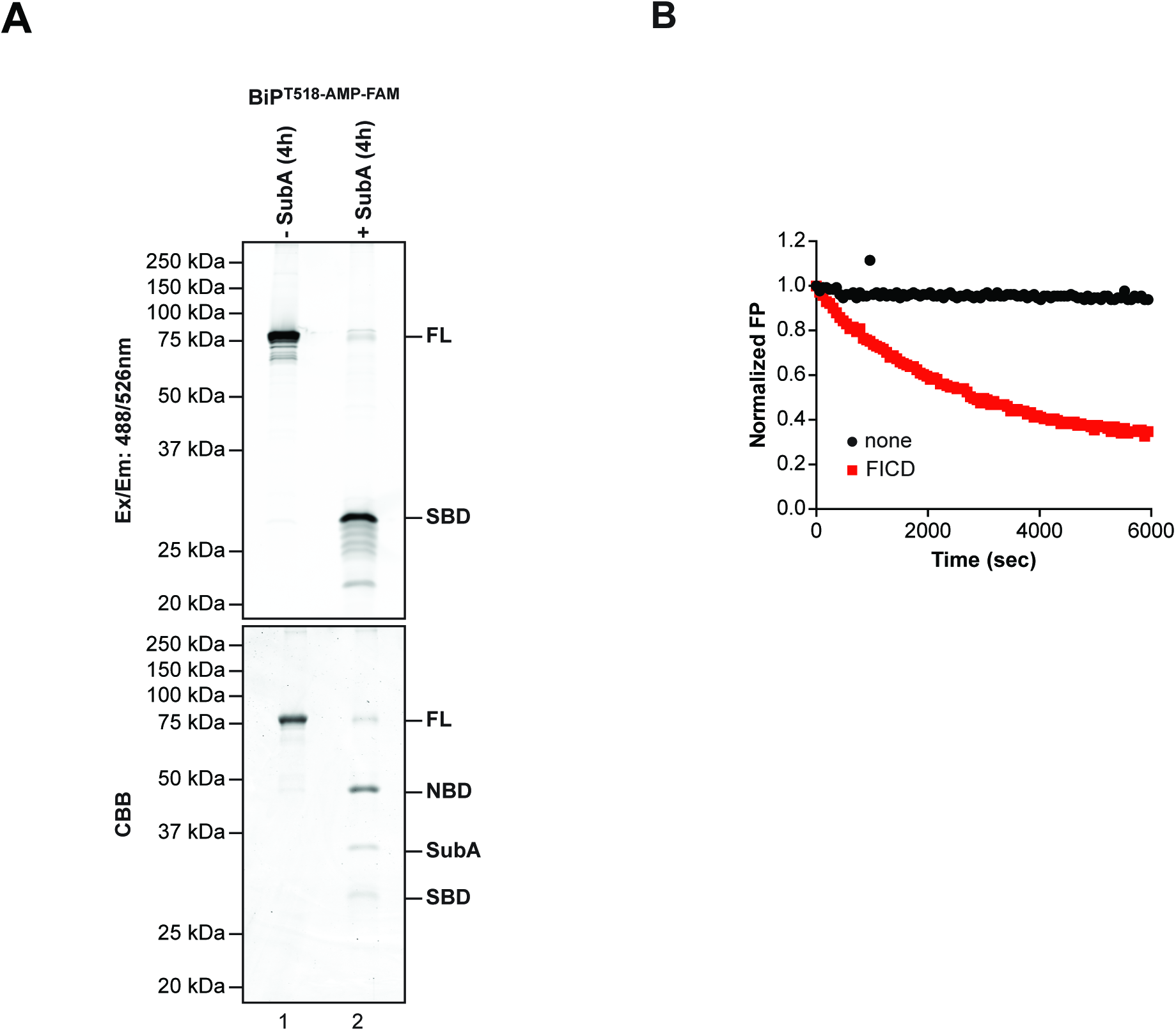
AMPylation with ATP-FAM generates BiP specifically labeled on its substrate binding domain. (**A**) SDS-PAGE gel of AMPylated BiP incubated without or with SubA protease. BiP was AMPylated in vitro with FICD^E234G^ in presence of a fluorescently labeled ATP derivative (ATP-FAM) and the resulting AMPylated BiP, with a fluorescent AMP attached (BiP^T518-AMP-FAM^) was re-purified (as in Figure 1D). After prolonged treatment without or with SubA (4 hours at 30°C) the samples were denatured and applied to SDS-PAGE. The fluorescence signals of the fluorophore in the gel were detected (excitation: 488 nm, emission: 526 nm; upper panel) and the proteins were visualized by staining with Coomassie (CBB; lower panel). Uncleaved full-length BiP^T518-AMP-FAM^ (FL), the nucleotide binding domain (NBD), the substrate binding domain (SBD), FICD, and SubA are indicated. (**B**) Time-dependent plot of fluorescence polarization (FP) of BiP AMPylated with FAM-labeled AMP (BiP^T518-AMP-FAM^, from the sample shown in “A” above) after incubation without (black line) or with wildtype FICD protein (red line). The decrease in the FP signal reflects release of the fluorophore from BiP.

**Figure 3 supplement 1.**
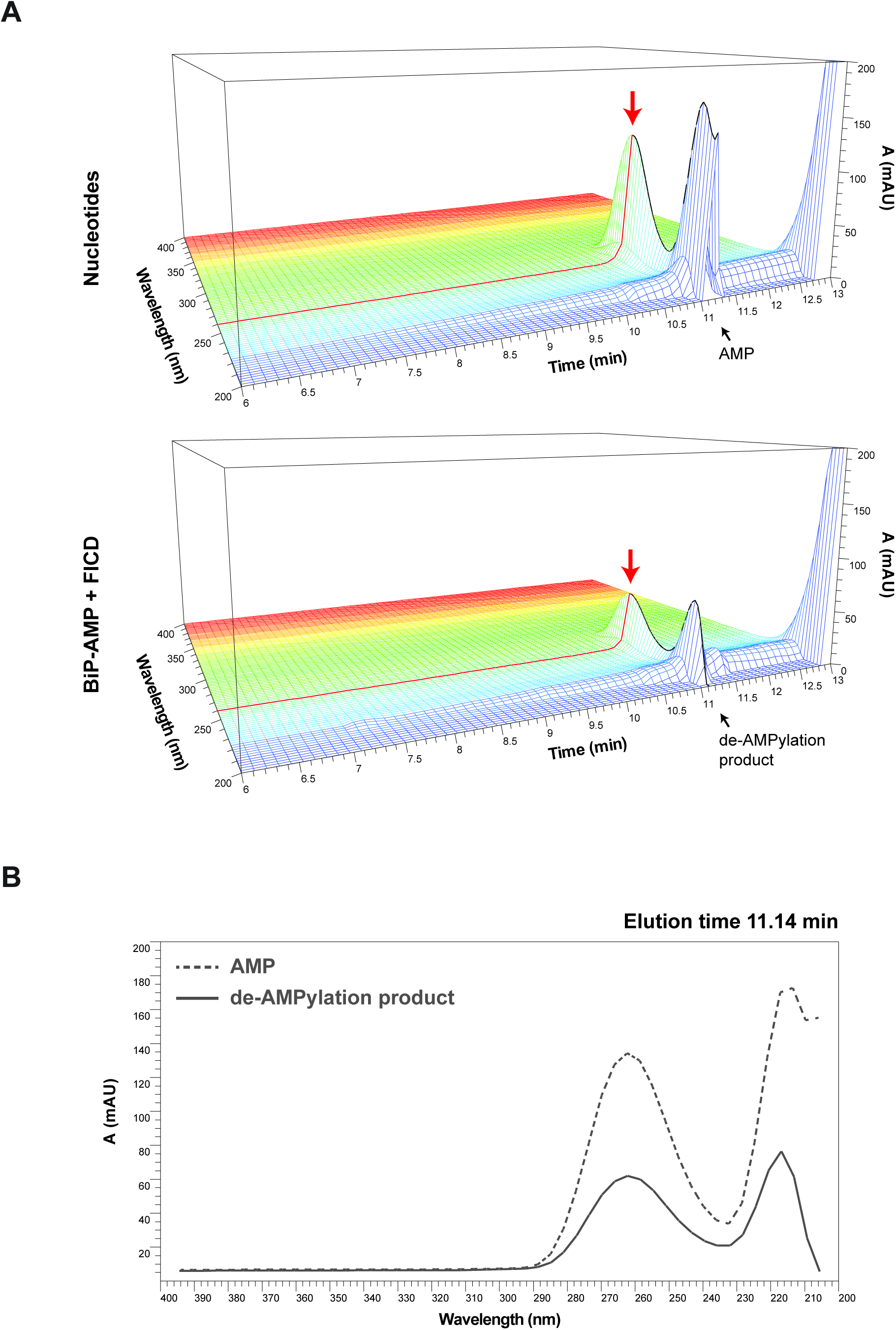
Absorbance spectra of ion pair chromatography elution profiles. (A) 3D absorbance plots of the nucleotide standard (upper panel) and the ‘BiPAMP+FICD’ de-AMPylation sample (lower panel) shown in Figure 4C (grey and red traces therein). The elution profiles between 6 and 13 minutes are shown. Note that the absorbance characteristics of the AMP standard and the de-AMPylation product (both eluting at ∼11.1 minutes) are qualitatively indistinguishable with an absorbance maximum at ∼260 nm (arrows). (B) Direct comparison of the absorbance spectra at 11.14 minutes of the profiles shown in “A”.

**Figure 6 supplement 1.**
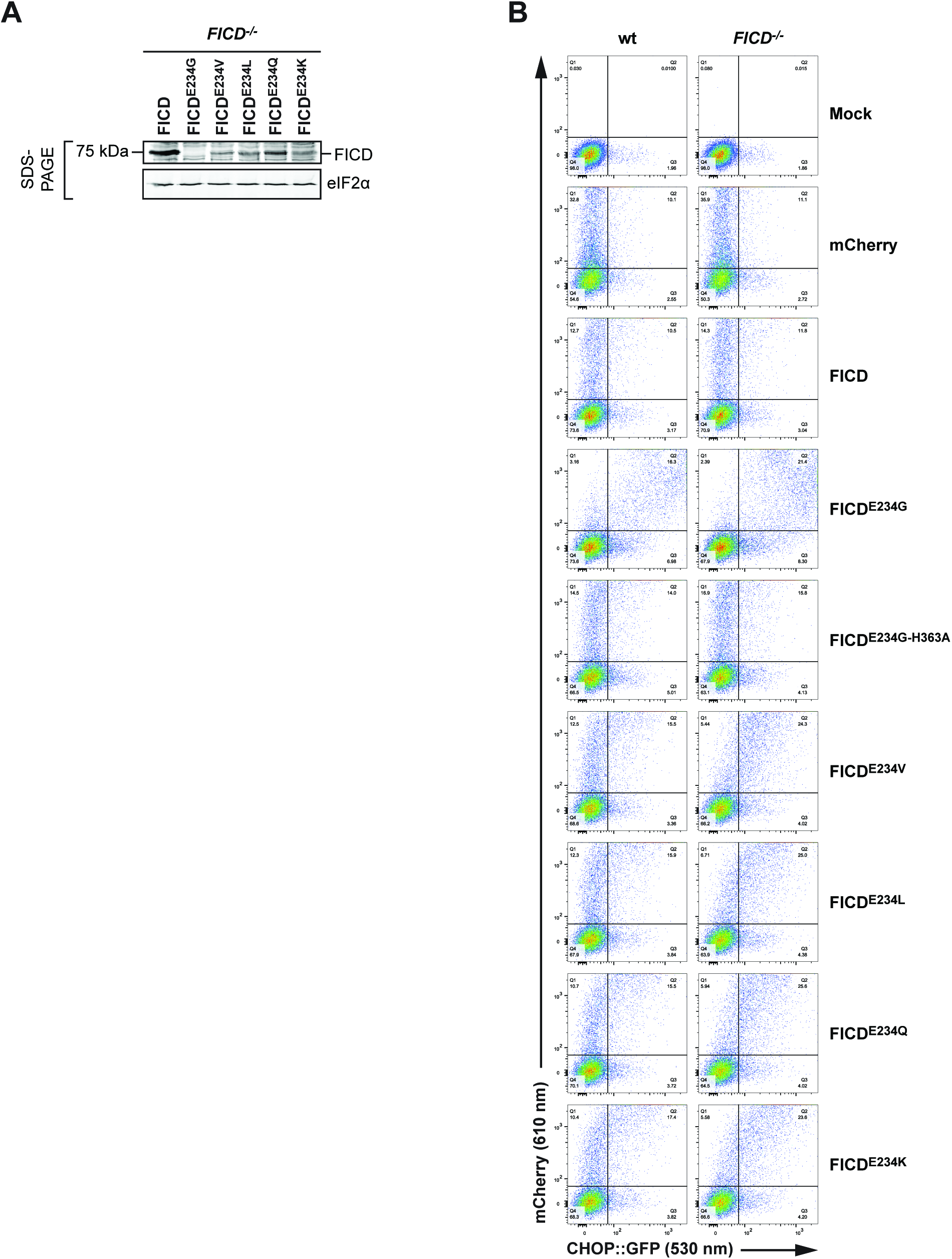
(A) FICD immunoblot of FICD-deficient (-/-) CHO-K1 cells transfected with plasmids encoding the indicated FICD derivatives. The eIF2α below serves as a loading control. Note the higher protein levels of the weaker AMPylation active/de-AMPylation defective FICD^E234V/L/Q/K^ mutants compared to the AMPylation hyperactive FICD^E234G^. (B) Flow cytometry source data from a representative experiment (one of three) used to generate the plot in Figure 6B.

**Figure 6 supplement 2.**
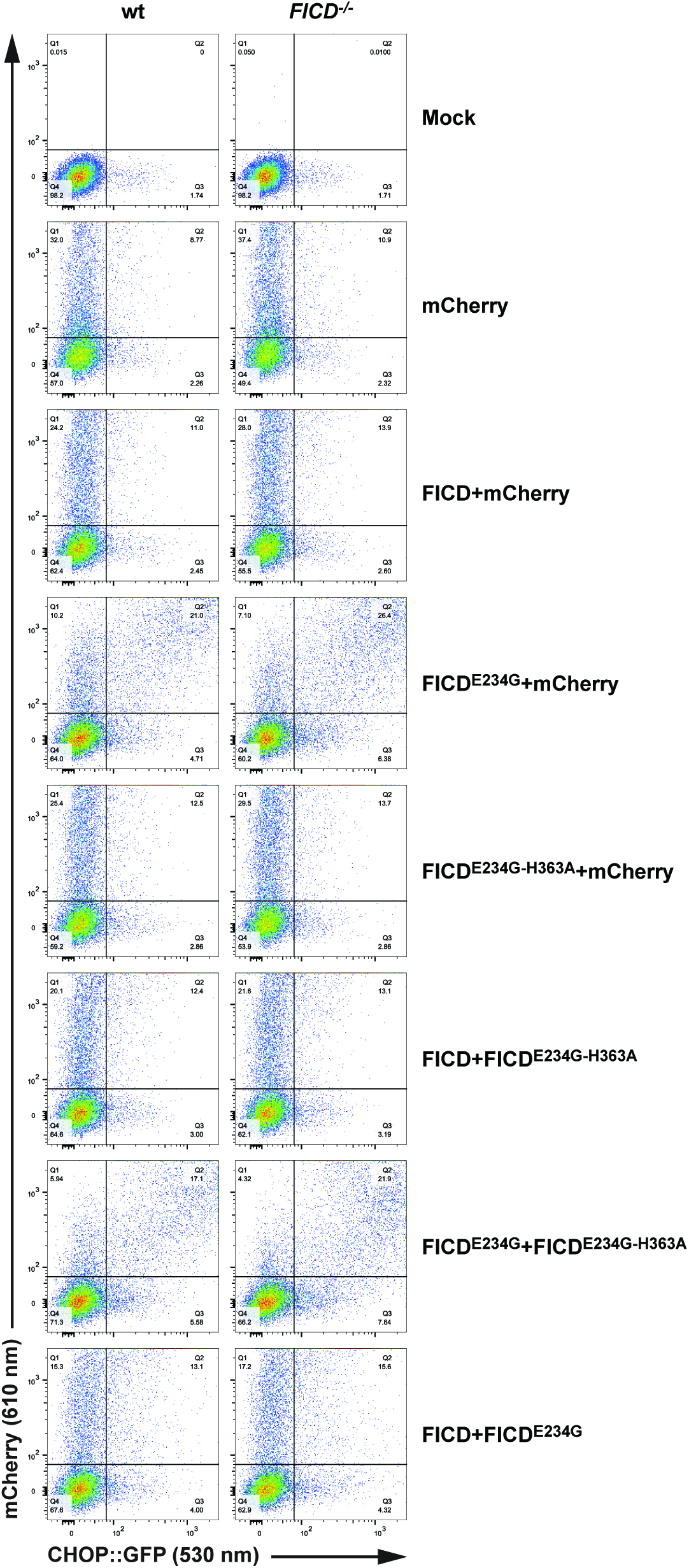
Flow cytometry source data from one of three independent repeats plotted in Figure 6C.

**Figure 6 supplement 3.**
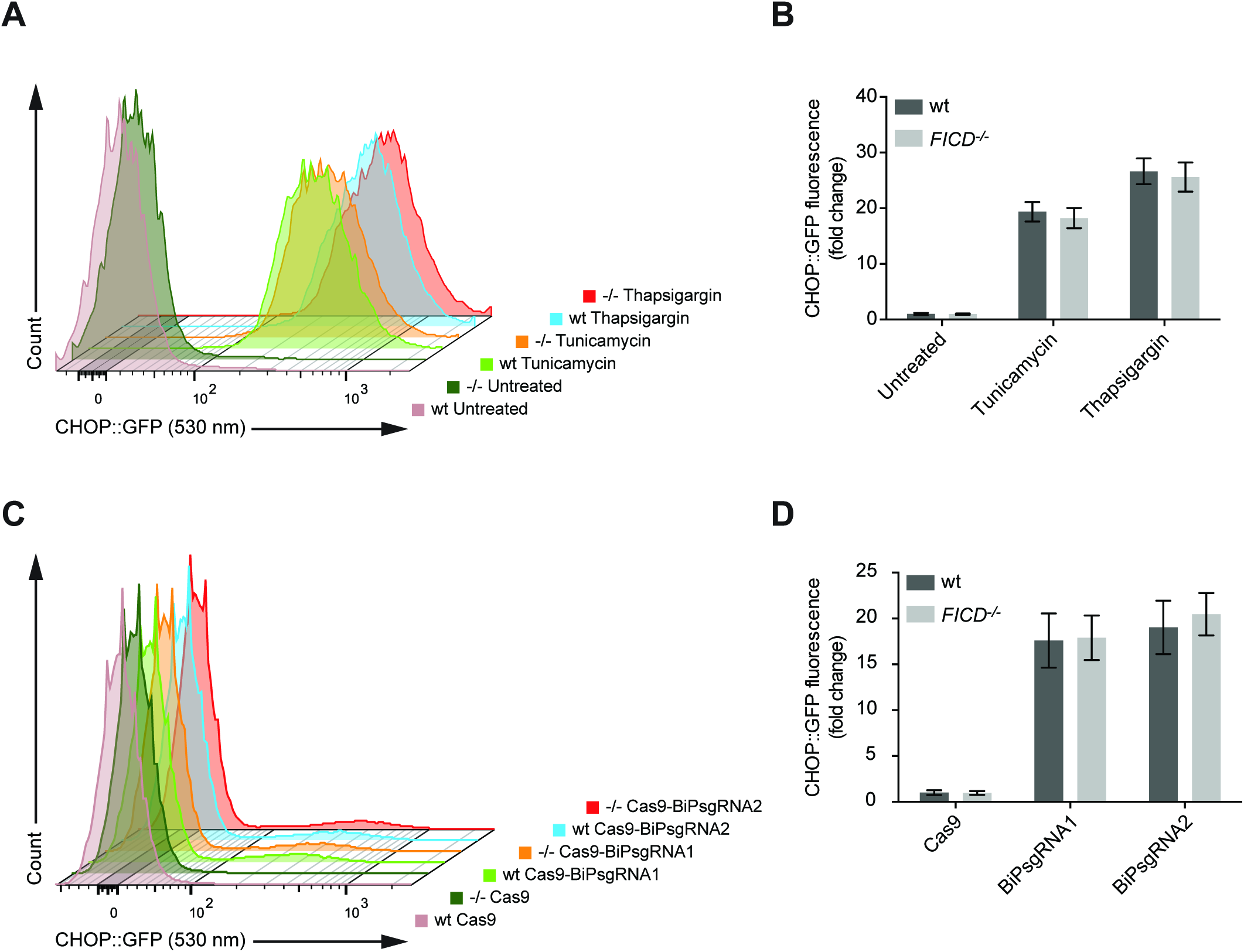
Wildtype and FICD-deficient (-/-) cells respond indistinguishably to unfolded protein stress in the ER. (A) Flow cytometry analysis of wildtype and FICD-deficient (-/-) CHO-K1 CHOP::GFP UPR reporter cells treated with the UPR-inducing compounds, tunicamycin (2.5 µg/ml) or thapsigargin (0.5 µM), for 16 hours before analysis. Note the equal accumulation of CHOP::GFP-positive cells in tunicamycin- or thapsigargin-treated wildtype and *FICD*^−/−^ cells. (B) Plot of the median values ± SD of the GFP fluorescent signal of the samples described in “A” from three independent experiments (fold change relative to untreated wildtype cells; *n* = 3). (C) Flow cytometry analysis of wildtype and FICD-deficient (-/-) CHO-K1 CHOP::GFP UPR reporter cells transiently transfected with plasmids encoding the Cas9 nuclease and single guide RNAs targeting hamster BiP. Note the similar levels of UPR signaling in wildtype and *FICD*^−/−^ cells. D) Plot of the median values ± SD of the CHOP::GFP fluorescent signal in the transfected subpopulation of the cells shown in “C” from three independent experiments. Transfected cells were identified by co-expression of a mCherry marker (not shown) carried by the Cas9 plasmid (fold change relative to wildtype cells transfected with plasmid DNA encoding Cas9 and mCherry only; *n* = 3).

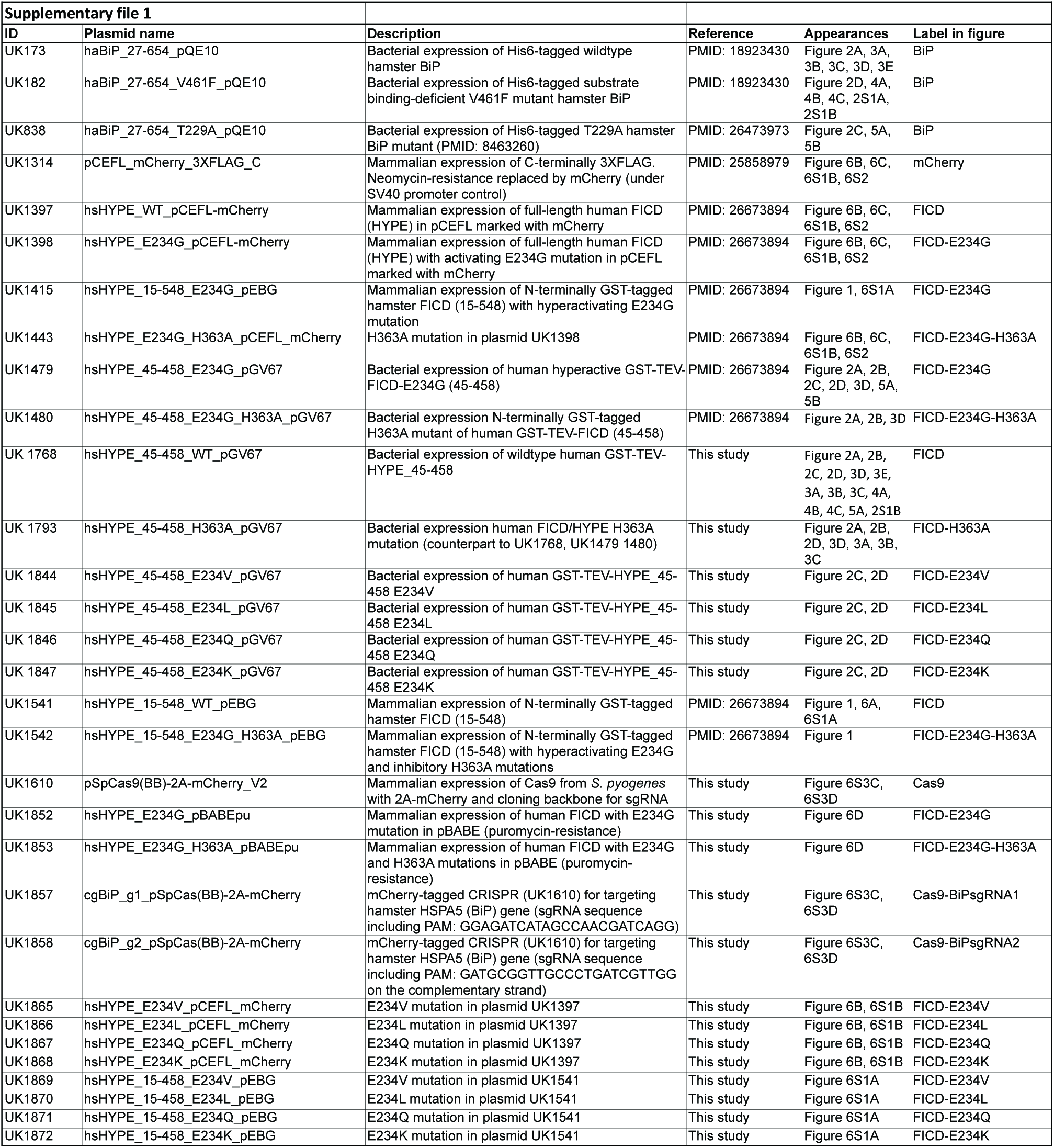

## References

Avezov, E., Cross, B.C., Kaminski Schierle, G.S., Winters, M., Harding, H.P., Melo, E.P., Kaminski, C.F., and Ron, D. (2013). Lifetime imaging of a fluorescent protein sensor reveals surprising stability of ER thiol redox. J Cell Biol 201, 337–349.

Balch, W.E., Morimoto, R.I., Dillin, A., and Kelly, J.W. (2008). Adapting proteostasis for disease intervention. Science 319, 916–919.

Bertolotti, A., Zhang, Y., Hendershot, L., Harding, H., and Ron, D. (2000). Dynamic interaction of BiP and the ER stress transducers in the unfolded protein response. Nat Cell Biol 2, 326–332.

Broncel, M., Serwa, R.A., Bunney, T.D., Katan, M., and Tate, E.W. (2015). Global profiling of HYPE mediated AMPylation through a chemical proteomic approach. Mol Cell Proteomics.

Bunney, T.D., Cole, A.R., Broncel, M., Esposito, D., Tate, E.W., and Katan, M. (2014). Crystal structure of the human, FIC-domain containing protein HYPE and implications for its functions. Structure 22, 1831–1843.

Carlsson, L., and Lazarides, E. (1983). ADP-ribosylation of the Mr 83,000 stress-inducible and glucose-regulated protein in avian and mammalian cells: modulation by heat shock and glucose starvation. Proc Natl Acad Sci U S A 80, 4664–4668.

Castro-Roa, D., Garcia-Pino, A., De Gieter, S., van Nuland, N.A., Loris, R., and Zenkin, N. (2013). The Fic protein Doc uses an inverted substrate to phosphorylate and inactivate EF-Tu. Nat Chem Biol 9, 811–817.

Chambers, J.E., Petrova, K., Tomba, G., Vendruscolo, M., and Ron, D. (2012). ADP ribosylation adapts an ER chaperone response to short-term fluctuations in unfolded protein load. J Cell Biol 198, 371–385.

Chevalier, M., King, L., Wang, C., Gething, M.J., Elguindi, E., and Blond, S.Y. (1998). Substrate binding induces depolymerization of the C-terminal peptide binding domain of murine GRP78/BiP. J Biol Chem 273, 26827–26835.

Engel, P., Goepfert, A., Stanger, F.V., Harms, A., Schmidt, A., Schirmer, T., and Dehio, C. (2012). Adenylylation control by intra- or intermolecular active-site obstruction in Fic proteins. Nature 482, 107–110.

Freiden, P.J., Gaut, J.R., and Hendershot, L.M. (1992). Interconversion of three differentially modified and assembled forms of BiP. EMBO J 11, 63–70.

Garcia-Pino, A., Zenkin, N., and Loris, R. (2014). The many faces of Fic: structural and functional aspects of Fic enzymes. Trends Biochem Sci 39, 121–129.

Gaut, J.R., and Hendershot, L.M. (1993). Mutations within the nucleotide binding site of immunoglobulin-binding protein inhibit ATPase activity and interfere with release of immunoglobulin heavy chain. J Biol Chem 268, 7248–7255.

Ham, H., Woolery, A.R., Tracy, C., Stenesen, D., Kramer, H., and Orth, K. (2014). Unfolded protein response-regulated Drosophila Fic (dFic) protein reversibly AMPylates BiP chaperone during endoplasmic reticulum homeostasis. J Biol Chem 289, 36059–36069.

Harms, A., Stanger, F.V., and Dehio, C. (2016). Biological Diversity and Molecular Plasticity of FIC Domain Proteins. Annu Rev Microbiol.

Khater, S., and Mohanty, D. (2015). In silico identification of AMPylating enzymes and study of their divergent evolution. Sci Rep 5, 10804.

Laitusis, A.L., Brostrom, M.A., and Brostrom, C.O. (1999). The dynamic role of GRP78/BiP in the coordination of mRNA translation with protein processing. J Biol Chem 274, 486–493.

Luong, P., Kinch, L.N., Brautigam, C.A., Grishin, N.V., Tomchick, D.R., and Orth, K. (2010). Kinetic and structural insights into the mechanism of AMPylation by VopS Fic domain. J Biol Chem 285, 20155–20163.

Michaelis, L., Menten, M.L., Johnson, K.A., and Goody, R.S. (2011). The original Michaelis constant: translation of the 1913 Michaelis-Menten paper. Biochemistry 50, 8264–8269.

Morgenstern, J.P., and Land, H. (1990). Advanced mammalian gene transfer: high titre retroviral vectors with multiple drug selection markers and a complementary helper-free packaging cell line. Nucleic Acids Res 18, 3587–3596.

Paton, A.W., Beddoe, T., Thorpe, C.M., Whisstock, J.C., Wilce, M.C., Rossjohn, J., Talbot, U.M., and Paton, J.C. (2006). AB5 subtilase cytotoxin inactivates the endoplasmic reticulum chaperone BiP. Nature 443, 548–552.

Petrova, K., Oyadomari, S., Hendershot, L.M., and Ron, D. (2008). Regulated association of misfolded endoplasmic reticulum lumenal proteins with P58/DNAJc3. EMBO J 27, 2862–2872.

Pincus, D., Chevalier, M.W., Aragon, T., van Anken, E., Vidal, S.E., El-Samad, H., and Walter, P. (2010). BiP binding to the ER-stress sensor Ire1 tunes the homeostatic behavior of the unfolded protein response. PLoS Biol 8, e1000415.

Preissler, S., Chambers, J.E., Crespillo-Casado, A., Avezov, E., Miranda, E., Perez, J., Hendershot, L.M., Harding, H.P., and Ron, D. (2015a). Physiological modulation of BiP activity by *trans*-protomer engagement of the interdomain linker. Elife 4, e08961.

Preissler, S., Rato, C., Chen, R., Antrobus, R., Ding, S., Fearnley, I.M., and Ron, D. (2015b). AMPylation matches BiP activity to client protein load in the endoplasmic reticulum. Elife 4, e12621.

Ran, F.A., Hsu, P.D., Wright, J., Agarwala, V., Scott, D.A., and Zhang, F. (2013). Genome engineering using the CRISPR-Cas9 system. Nat Protoc 8, 2281–2308.

Ronda, C., Pedersen, L.E., Hansen, H.G., Kallehauge, T.B., Betenbaugh, M.J., Nielsen, A.T., and Kildegaard, H.F. (2014). Accelerating genome editing in CHO cells using CRISPR Cas9 and CRISPy, a web-based target finding tool. Biotechnol Bioeng 111, 1604–1616.

Roy, C.R., and Cherfils, J. (2015). Structure and function of Fic proteins. Nat Rev Microbiol 13, 631–640.

Sanyal, A., Chen, A.J., Nakayasu, E.S., Lazar, C.S., Zbornik, E.A., Worby, C.A., Koller, A., and Mattoo, S. (2015). A novel link between Fic (filamentation induced by cAMP)-mediated adenylylation/AMPylation and the unfolded protein response. J Biol Chem 290, 8482–8499.

Scorsone, K.A., Panniers, R., Rowlands, A.G., and Henshaw, E.C. (1987). Phosphorylation of eukaryotic initiation factor 2 during physiological stresses which affect protein synthesis. J Biol Chem 262, 14538–14543.

Walter, P., and Ron, D. (2011). The unfolded protein response: from stress pathway to homeostatic regulation. Science 334, 1081–1086.

Wang, M., and Kaufman, R.J. (2016). Protein misfolding in the endoplasmic reticulum as a conduit to human disease. Nature 529, 326–335.

Worby, C.A., Mattoo, S., Kruger, R.P., Corbeil, L.B., Koller, A., Mendez, J.C., Zekarias, B., Lazar, C., and Dixon, J.E. (2009). The fic domain: regulation of cell signaling by adenylylation. Mol Cell 34, 93–103.

Xiao, J., Worby, C.A., Mattoo, S., Sankaran, B., and Dixon, J.E. (2010). Structural basis of Fic-mediated adenylylation. Nat Struct Mol Biol 17, 1004–1010.

Xu, Y., Carr, P.D., Vasudevan, S.G., and Ollis, D.L. (2010). Structure of the adenylylation domain of E. coli glutamine synthetase adenylyl transferase: evidence for gene duplication and evolution of a new active site. J Mol Biol 396, 773–784.

